# Cancer cell adaptations along a proliferation–robustness axis drive TME pro-tumor phenotypes

**DOI:** 10.64898/2026.10.27.762245

**Authors:** Maayan Pour, Guisheng Zhao, Angeliki Karatza, Gustavo S. França, Jingye Wang, Linda Procell, Anjali Rao, Deborah A. Corcoran, Ayushi S. Patel, Cristina H. Hajdu, Kwok-Kin Wong, Amanda W. Lund, Edna Cukierman, Itai Yanai

## Abstract

Tumors are highly complex dynamic ecosystems with causal relationships between cancer cells and their tumor microenvironment (TME). To disentangle these interactions, we performed long-term co-culture experiments with cancer cells evolving alongside continuously replenished fibroblasts or macrophages. This allowed us to distinguish between adaptive and physiological responses in specific cancer-TME interactions, which we then also recapitulated in patient tumors. We found that co-culture with fibroblasts induced the epithelial-mesenchymal transition in cancer cells and resulted in their increased drug tolerance, while co-culture with macrophages promoted stress, hypoxia responses and enhanced tumorigenicity. Adaptations were recurrent and partially reversible, but could not be explained by clonal selection, suggesting a dynamic non-mutational process. Moreover, adapted cancer cells elicited cell state changes in the fibroblasts and macrophages consistent with an “extended phenotype” mode of cancer evolution. Our findings reframe cancer-TME interactions by a systematic dissection according to sets of specific adaptations and programmed responses.

## INTRODUCTION

A malignant tumor comprises a complex network of evolving interactions among its numerous cellular components, including cancer, fibroblast, endothelial, immune and other cell types^1,2^. As a reflection of such interactions, malignant cells exhibit diverse cellular states across cancer types. These cellular states have been defined mostly on the basis of gene expression^3–10^ and include aspects of both biological processes – such as stress, interferon-response, and hypoxia – and cellular identity-like states – including AT2-like and basal-like in lung cancer^4,10^. The recurrence of these states across tumors highlights their central role in tumorigenesis and disease outcomes. Notably, the EMT (or EMT-like) cell state has been associated with metastasis and drug resistance in several cancer types^11^ and an hypoxic state has been associated with immunosuppression in glioblastoma^12^. But while principles regarding the developmental constraints on the cellular identity states are beginning to emerge^13^, and the possibility of cooperation among cells has also been evidenced^14^, a conceptual framework to explain the composition and dynamics of cancer cell states remains elusive.

Diverse cellular states have also been identified within the non-malignant compartments of the TME, linked with patient prognosis^15^. Cancer-associated fibroblasts (CAFs) often diversify into many functional states, such as antigen-presenting, myofibroblastic, or inflammatory, which were originally characterized in wound healing needed functions, and range from anti-tumoral to pro-tumoral modalities^10,16–25^. Similarly, tumor-associated macrophages (TAMs) can adopt inflammatory states with anti-tumoral properties (such as IFN-stimulated macrophages) or anti-inflammatory states with immunosuppressive functions, akin to macrophages partaking action in chronic wound healing^26–28^. T-cell subtypes also differ in their roles, with cytotoxic T cells exhibiting anti-tumoral features, whereas regulatory T cells (Tregs) promote tumor progression^29^. Finally, evidence indicates that cancer cell states are spatially organized relative to the TME cell types and exhibit specific dependencies that establish interaction niches within the TME^2,10,30,31^.

When considering interactions between cancer and TME cells, we reasoned that it is important to distinguish between two modes of interactions. First, upon initial contact, the cancer and TME cells may each trigger a *physiological* response to the presence of the other; in other words, a reciprocal programmed cell state change. For example, fibroblasts may be induced to activate various transcriptional programs following biochemical and mechanical force changes of the extracellular matrix (ECM), such as increased tissue stiffness^32^. Second, as a consequence of natural selection, *adaptation* may occur such that the cellular states would correspond to evolved responses^33^, unfolding over the long-term course of the interaction. This second mode – which may occur by genetic (mutations) or non-genetic (plasticity) mechanisms – carries important implications for how we view these interactions and their consequences for tumorigenesis, yet they have not been systematically studied.

Here, we designed an experiment to test whether extended interactions between cancer cells and cell types from the TME can elicit adaptations in the cancer cells. By carrying out a long-term co-culture, in which cancer cells evolved in the presence of TME cell types that were replenished weekly, we were able to distinguish between an immediate response to an interaction – which remains constant over time (that is, physiologically-induced) – and those developing over many weeks (that is, adaptive). When studying patient samples, we found evidence for corresponding adaptations observed *in vitro*, suggesting that this approach can parse out distinct behaviors within the complex tumor ecosystem. We also revealed that adaptive responses exhibit signatures of recurrence, reversibility, and generality across cancer types. Interestingly, adaptive cell state changes in cancer cells were specific to their co-cultured TME-cell types, thus revealing a functional trade-off. Finally, we found that the adapted cancer cells were able to elicit distinct states in the TME cells over time, providing evidence that tumor-associated states can be considered an extended phenotype of the cancer. Functionally testing this relationship, we observed that disrupting a macrophage-adapted program in the cancer cells resulted in a reversion in the macrophage state. Our approach demonstrates the power of studying adaptive cancer processes using simple systems by isolating specific dynamic cancer-TME interactions that occur in the highly complex tumor ecosystem.

## RESULTS

### Long-term co-culture of cancer cells with replenished distinct TME cell types leads to phenotypic adaptations

Cancer cell states tend to be associated with distinct compartments of the tumor ecosystem^10,16–19,22,34–36^, leading us to ask whether the association between cancer cell states and microenvironments reflect (1) immediate responses to that microenvironment, (2) beneficial adaptations through interactions with the environment over time, or (3) a combination of these two. To distinguish among these possibilities we devised a long-term co-culture system, whereby cancer cells are allowed to evolve over 36 weeks in the presence of primary non-cancerous cell types (Fig. 1A). As we co-cultured primary normal cell types with cancer cells, we henceforth refer to them as TME cells. By replenishing these cells weekly with fresh naive murine fibroblasts or macrophages, we reasoned that this long-term experiment would allow us to distinguish between physiological and adaptive responses of the cancer cells. Importantly, because this experiment exclusively follows the evolution of the cancer cells, it is designed to test whether such co-cultured cancer cells can reconstitute both the cancer and TME states observed in tumors.

**Figure 1.**
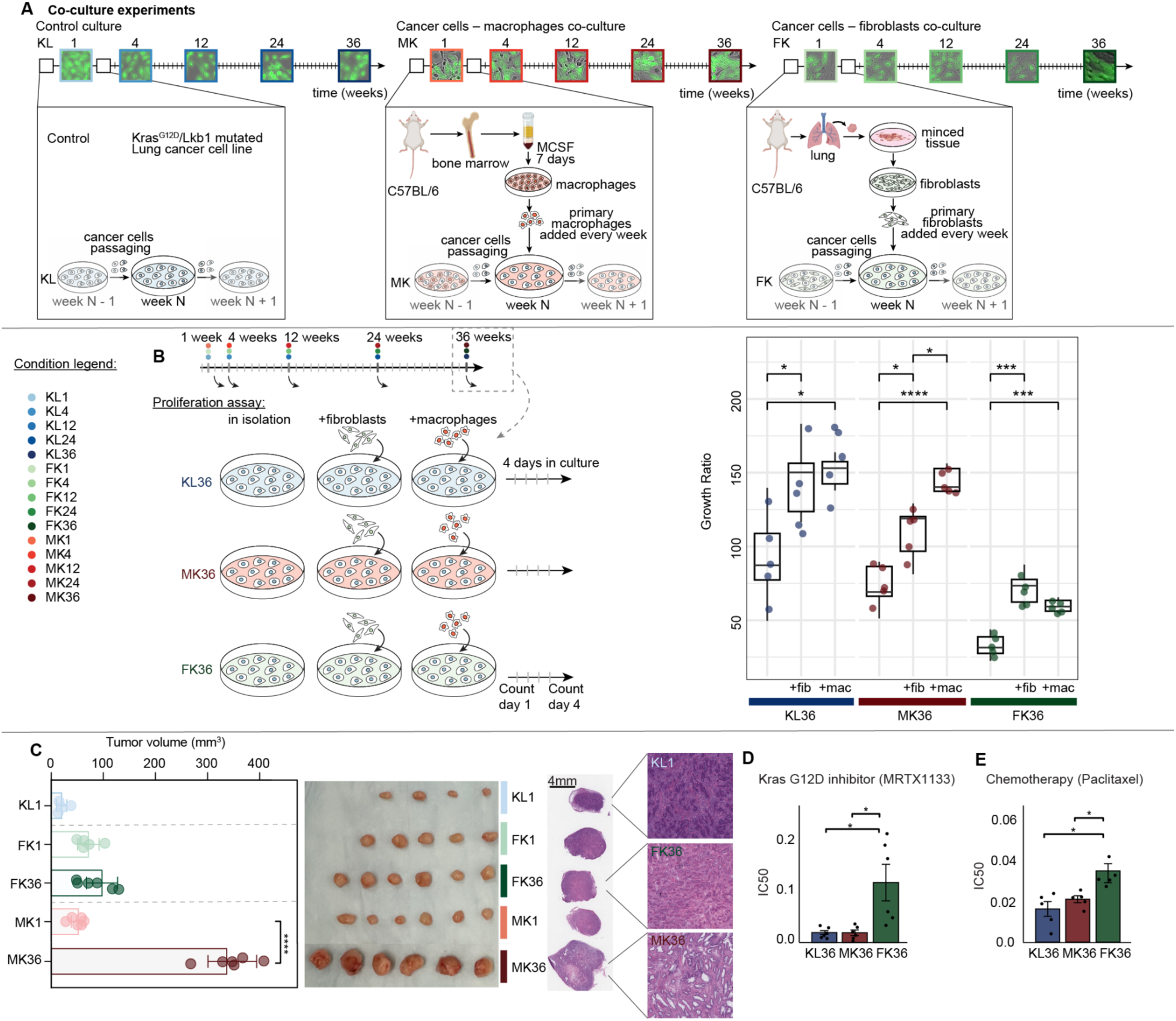
Long term *in vitro* co-culture of cancer cells with replenished macrophages or fibroblasts leads to phenotypic changes. **A.** Experimental design of the long-term co-culture experiment. The KLa lung cancer cell line (*Kras*^G12D^/*Lkb1*^-/-^mutated) was cultured for 36 weeks under three conditions: (1) control, no co-culture (KL), (2) with freshly added primary macrophages (MK), and (3) with freshly added fibroblasts (FK). In each week of the experiment, the cancer cells of the co-culture were re-plated in a new dish, with macrophages or fibroblasts, freshly isolated from mice weekly (see Methods). In the experiment timeline, ticks represent the week of replating and boxes show micrographs (scale, 100μm) representing cells collected for single-cell RNA-seq and downstream experiments. As a marker of the cancer cells relative to the fibroblasts and macrophages, we added GFP to the ancestral line prior to initiating the co-cultures. **B.** Schematic (left) and box plot (right) of the proliferation assay of the KL_36_, FK_36_ and MK_36_ lines in isolation, and together with fibroblasts/macrophages. All experiments were performed in 5 technical replicates. **C.** Tumor volume measurements 32 days following subcutaneous injection of the co-culture cell lines in mice. 4-6 tumors were collected from 2-3 mice for each condition. The tumors from the MK_36_ line were significantly larger than all the rest (*P* < 10^-4^). The tumors from FK_36_ lines were significantly larger than the KL_1_ lines (*P* < 10^-2^). The rest were not significantly different. H&E images from representative tumors. **D, E.** Cell viability of the 36-week co-cultured cancer cell lines during 3 days of treatment with G12D inhibitor (**D**) or chemotherapy (**E**). IC50 bar plots represent the average 6 technical replicates per experiment with their standard deviations. Values were normalized relative to vehicle-treated controls for each respective cell line. As these two drugs elicit distinct responses the detected sensitivity is shown on a different scale. Significance calculated by t-test. * *P* < 0.05, ** *P* < 10^-2^, *** *P* < 10^-3^, **** *P* < 10^-4^

As our main model co-culture system, we adopted the KLa mouse lung cancer cell line (*Kras*^G12D^/*Lkb1*^-/-^mutant^37^). We also tested for the generality of our results using additional cell lines (see below). To isolate and study individual interactions observed in the tumor architecture, we carried out co-cultures of cancer cells with two key cell types of the TME cellular components: macrophages in one arm of the experiment (MK), fibroblasts in another (FK), and as a control, cancer cells maintained in isolation (KL) (Fig. 1A). We performed these experiments in independent quadruplicates for a total of 36 weeks. Throughout the experiment, we collected cells after 1, 4, 12, 24 and 36 weeks to study their phenotypic and gene expression changes. This resulted in the MK_1_, MK_4_, MK_12_, MK_24_, and MK_36_ cancer cell lines from the co-culture with macrophages; the FK_1_, FK_4_, FK_12_, FK_24_, and FK_36_ cancer cell lines from the co-culture with fibroblasts, and the same for the KL samples for the control non-co-cultured lines.

These names relate to the cancer cells that were retrieved from the co-culture. We also collected macrophages and fibroblasts from the co-culture throughout the different time-points. We named them explicitly ‘macrophages from co-culture with:’ and ‘fibroblasts from co-culture with:’ alongside the cancer cells they were cultured with. In each week of the experiment, the cancer cells were passaged and maintained in the co-culture, while the TME cell types were added fresh from primary cultures, isolated from mouse bone marrow or lung, at a ratio of 1:50 (cancer cells to macrophages) for the MK and 1:10 (cancer cells to fibroblasts) for the FK lines (Fig. 1A). To eliminate initial heterogeneities, we initiated the experiment after expanding a single clone from the KLa line.

If adaptation occurred over the course of the co-culture, we would expect that, in the presence of the TME cell type of the co-culture, a cancer cells’ fitness advantage would occur. We thus performed a live imaging-based proliferation assay, to test for increased fitness among the resulting KL_36_, MK_36,_ and FK_36_ co-cultured lines; in other words, those cancer lines co-cultured for 36 weeks. We tested the proliferation rates of the KL_36_, MK_36,_ and FK_36_ lines (cancer cells only), both in isolation and co-cultured with fibroblasts or macrophages, over 4 days (Fig. 1B and Fig. S1). We found that in the presence of either fibroblasts or macrophages, proliferation rates increased in all three lines (KL_36_, MK_36_ and FK_36_, Fig. 1B). This is consistent with previous work showing that addition of fibroblasts and macrophages leads to enhanced proliferation^38,39^. However, in the MK_36_ and FK_36_ lines we found TME cell-type specific effects: each exhibited increased proliferation in the presence of the cell type with which they had been co-cultured (Fig. 1B; though this change is not significant for FK_36_). Most notably, adding macrophages to MK_36_ cells led to a nearly 3-fold increase in their proliferation (*P* < 0.05), while adding fibroblasts to this line only led to a 2-fold increase. Testing the proliferation rates of the KL_1_, MK_1,_ and FK_1_ lines – those lines co-cultured for only 1 week – in the presence of either fibroblasts or macrophages, revealed similar proliferation rates of the cancer cells, suggesting that the differences between the lines observed in the 36-week cell lines are an adaptive effect, as opposed to an immediate one (Fig. S1A). Indeed, we observed that in the KL_12_, MK_12_, and FK_12_ lines (after 12 weeks of the co-culture experiment), the MK_12_ and FK_12_ lines showed a slight but non-significant preference for proliferation in the presence of macrophages and fibroblasts, respectively (Fig. S1B). The FK lines showed consistently slower baseline proliferation (for example, *P* < 0.05, FK_1_-MK_1_, Fig. S1).

In summary, we found evidence for adaptation in cancer lines continuously cultured with TME cell types by observing that: (1) while the addition of either cell type led to an enhanced proliferation rate in the control line (KL), the MK and FK lines exhibited a cell-type preference matching the cell type of their long-term co-culture (significantly only in the MK line), indicating an environment-specific proliferation advantage, and (2) the proliferation advantage increased over time (Fig. S1A, B), supporting the notion that this constitutes an adaptive process distinct from an immediate physiological (programmed) response upon co-culture.

To further test for the phenotypic consequences of the cancer’s TME adaptation, we generated subcutaneous tumors in mice and queried for their tumor growth capacity (Fig. 1C). Our proxy for a cell line’s inherent level of proliferation was the size of the tumor it spawns. We examined mice tumors generated by the MK_36_ lines and observed that the adaptation to macrophages led to significantly larger tumors relative to MK_1_ (*P*<10^-4^). In contrast, tumors generated using the FK_36_ line were not significantly different in size than those generated by the FK_1_ line (Fig. 1C). We further tested the effect of co-culture with macrophages over time on tumor growth, and found increasingly larger tumors along the adaptation and that these tumors had larger solid tumor cores and areas of necrosis, suggesting a more rapid tumor growth (Fig. S2A-B).

In addition to proliferation as a phenotypic consequence, we assessed whether long-term co-culture with the TME cell types leads to distinct survival capacities. We tested for drug tolerance by exposing the 36-week co-cultured cancer cell lines – KL_36_, FK_36_ and MK_36_ lines, without the addition of the TME cell types – to a Kras^G12D^ inhibitor (MRTX1133) and chemotherapy (Paclitaxel). After treating the lines for 3 days we then assessed their drug tolerance by measuring their IC50. Interestingly, we found that the cancer cells that had been co-cultured with fibroblasts showed significantly less sensitivity to these drugs (*P* < 0.05) (Fig. 1D-E, Fig. S2C). Consistently, tumors with higher stromal abundance have been shown to be less sensitive to KRAS^G12D^ inhibitors^40^. As a control, we also tested for the sensitivity to these drugs of cancer cells co-cultured for just one week and did not observe a significant difference between conditions (Fig. S2D), suggesting that the ability to better survive drug treatments emerged over the course of the long-term co-culture. Taken together, we found that long-term co-culture resulted in phenotypic changes that are environment-specific suggesting an adaptation that leads to a fitness advantage in that environment, and along a proliferation-robustness axis.

### Adaptive and physiological cancer cell state responses to TME interactions

The phenotypic changes observed in the co-cultured lines led us to investigate the molecular basis of this adaptation. We performed single-cell RNA sequencing (scRNA-Seq) to profile gene expression changes of the co-cultured cancer cells, for 5 representative stages throughout the experiment (Fig. 1A). The scRNA-Seq data allowed us to clearly separate between the cancer cells (KL_1-36_, MK_1-36_ and FK_1-36_), and the macrophages or fibroblasts extracted from the co-culture (Tables S8 and S9 and Figs. S3 and S16). When subjecting the three sets of cancer lines to dimensional reduction (KL, MK, and FK; and ensuring that only cancer cells are included in the analysis, Fig. S3), we observed that these cancer cell transcriptomes did not form distinct clusters (Fig. 2A). This suggested an induction of specific cell state changes, as opposed to large cell identity transformations. To identify the gene modules underlying these state changes, we used a gene module detection approach^41^ to systematically characterize the patterns of gene expression changes in the three lines over time. In contrast to a global expression analysis, detecting differentially expressed genes across conditions (Fig. S3D,E), the gene module detection approach does not rely on information of the cell labels, but rather generally identifies co-expressed genes. Applying this analysis, we delineated 15 modules, each with an average of 58 genes (Fig. 2B and Table S1). We detected similar gene modules using a non-negative factorization (NMF) approach^42^ (Table S2, Fig. S4E). To infer functional associations, we compared each module’s set of genes with previously defined cancer gene programs^4,10^ by gene set enrichment analysis^43,44^. This allowed us to relate the modules induced in the co-culture experiment to many previously observed modules in patient tumors, including EMT, secretion and stress^4,10^, suggesting the relevance of this *in vitro* system to *in vivo* processes (Fig. S4B, C).

**Figure 2.**
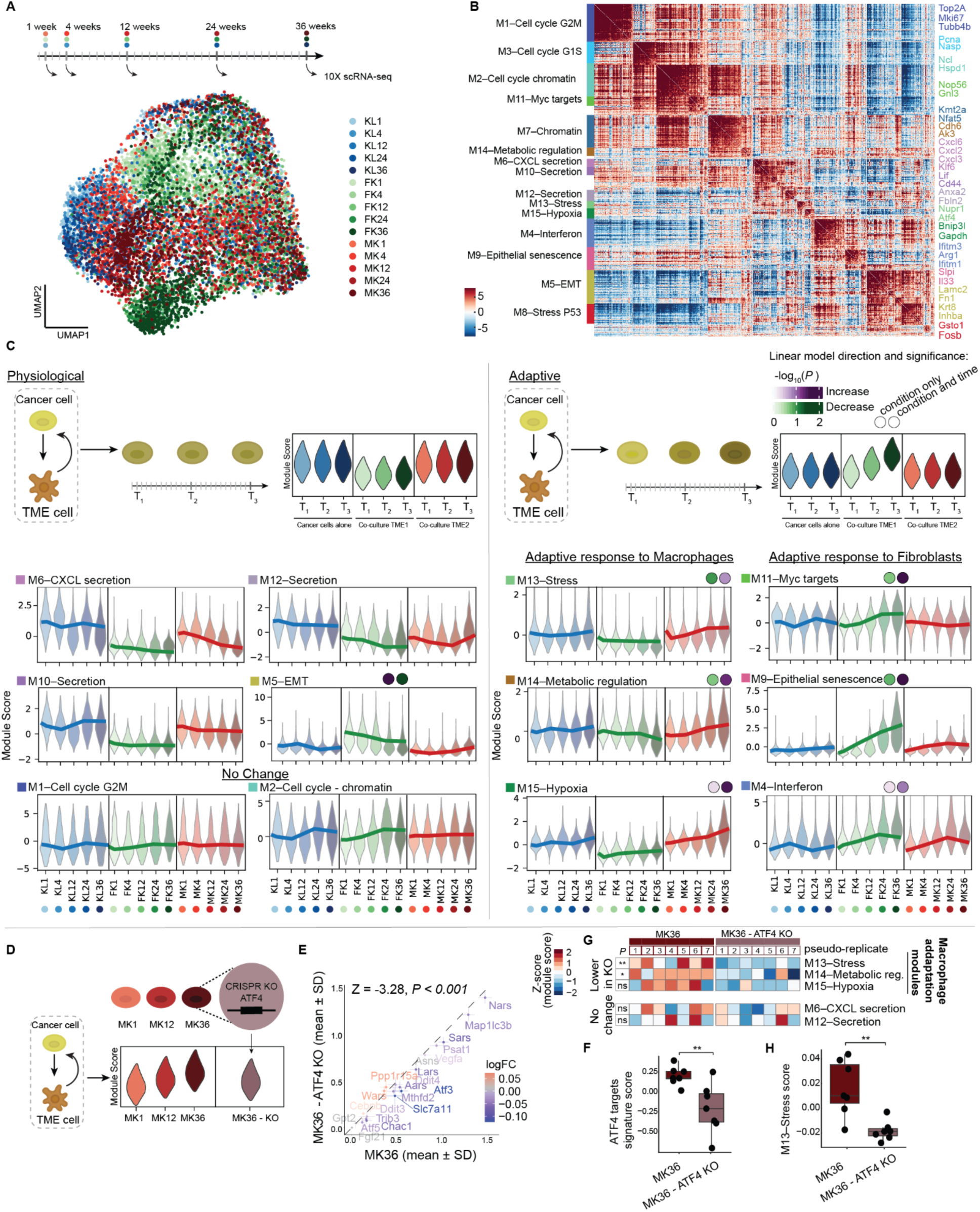
Gene modules reveal physiological and adaptative response programs in cancer cells co-cultured with TME cells. **A.** A UMAP projection of cancer cells from the co-culture experiment. Cells are colored by their condition and weeks in co-culture. **B.** Heatmap of gene-gene correlations (Z-scored), clustered as 15 modules detected by Hotspot analysis. The indicated genes are representatives from each module, and the modules were named based on classification of the genes to main functions and overlapping genes with previously defined modules (Fig. S5). **C.** Violin plots of the scores of the indicated Hotspot gene modules across cells from co-culture conditions and timepoints. (top) Schematic of the classification of modules by the score of their expression in the co-cultured cells. (left) Physiological response modules with a significant change due to the co-culture, with one or both of the cell lines, regardless of time in the co-culture. (right) Adaptation modules with a gradual change in the co-culture over time. For the adaptive modules, the *P* value of the linear model for change due to change of condition (physiological, left) or for the change over time (adaptive, right) is marked above the relevant condition (FK/MK). Purple marks increase and green marks decrease. **D.** Schematic of the module perturbation experiment. Cells were co-cultured with macrophages for 36 weeks, resulting in an increased expression of the M13–Stress module. CRISPR knockout of ATF4, a stress regulator from the M13 module, to test whether the expression of the module is dependent or not upon a common regulator. **E.** Plot showing the level of expression (mean and SD) of ATF4 targets^53^, in the MK_36_ cells versus the MK_36_ ATF4-KO cells. Z score and *P* value calculated against a random list of genes in the same length and from the same bins of expression levels. Cells were randomly assigned to seven pseudo-bulk groups to allow statistical testing. **F.** Boxplot comparison of the expression of the ATF4 targets as a signature, between MK_36_ cells and the MK_36_ ATF4-KO cells. **G.** Heatmap of averaged expression of macrophage adaptation modules and modules that were not affected by the ATF4 KO, across pseudo bulk cell groups of MK_36_ controls, and ATF4 knock out. *P* values were calculated using the Wilcoxon test for the differences between groups. **H.** Boxplot comparison of the expression of the M13–Stress module scores between MK_36_ cells and the MK_36_ ATF4-KO cells.

We sought to reveal for each module whether it captured a ‘physiological’ or an ‘adaptive’ response of the cancer cells to the presence of the TME cells. We expected that a module induced physiologically (as an immediate programmed response) would be present already in the first week of the co-culture and then remain constant throughout the remaining weeks; while an adaptive response module would have an induction that increased over time. We thus scored the cancer cells for the module expression and compared across the weeks of the co-culture (Fig. 2C, Fig. S4A). Using a linear model, we then calculated for each module, in each condition, the degree of change over time. For each module, the induction in the KL lines did not show a higher change over time compared to that in the MK and FK lines. Moreover, in the MK and FK lines, four and five modules significantly changed over time, respectively (Fig. S5).

Thus, despite minor changes due to long-term cell culture, most of the module variation over time reflected expression changes driven by co-culture with fibroblasts or macrophages. Based on our temporal score and the linear model significance, we then classified the modules into three broad categories: (1) physiological responses that occur immediately with the introduction of the TME cell types, (2) adaptive responses reflecting changes that unfold gradually over time in the co-culture with macrophages or fibroblasts, but not in the KL cells cultured alone and (3) cellular process independent of co-culture condition; indicating no significant change between conditions, or temporally within a co-culture condition (Fig. 2C, Fig. S5).

We detected strong physiological changes of expression that occurred immediately with the introduction of both fibroblasts and macrophages, though the effect was weaker with that of macrophages. We noted a decrease in expression of the M6–CXCL secretion and M12– Secretion in both co-cultures and, specifically in the fibroblast co-culture, an increase in M5– EMT and decrease in M10–Secretion (Fig. 2C, left). M6–CXCL secretion, M12–Secretion, and M10–Secretion are related to chemokine secretion and ECM remodeling. M6–CXCL was enriched with *CXCL* genes (e.g., *Cxcl1*, *Cxcl2*), which are known to be secreted to elicit immune cell recruitment^45–47^. *CXCL* upregulation is associated with inflammatory response and immune recruitment, which may be expected to decrease following TME interactions^48^. This process is part of a broader immunosuppression (tumor-promoting) mechanism that can involve downregulation of immune-stimulating molecules such as *CXCL*s while increasing the production of immunosuppressive factors such as *TGF-β*, *IL-10*, and other immune-modulatory cytokines^48^. M5–EMT included *Fn1* and *Krt18* suggesting that the presence of fibroblasts in the culture draws more cell adhesion changes, in particular, a significant increase in the EMT-like program^49–51^. Moreover, the EMT signature was highly expressed in all timepoints in the FK co-culture lines when compared to KL and MK lines, which may at least in part explain the slower proliferation of the adapted FK lines (Fig. 1B). Comparing across timepoints though, we also noted a decrease in the EMT module over time within the FK populations, which may indicate that the previously described^10,19^ spatial correlation in tumors between cancer cells in the EMT state and fibroblasts reflects an invasion of cancer cells into the stroma, rather than long-term proximity between those populations.

Unique to this experiment, we identified adaptive responses – modules of genes whose expression increases over time – following co-culture with fibroblasts (M4–interferon, M9– Epithelial senescence, and M11–Myc targets) or macrophages (M13–Stress, M14–Metabolic regulation, and M15–Hypoxia) (Fig. 2C, right). Inductions of these modules exhibited gradual expression changes in the cancer cells over time, suggesting that they emerged through a cancer-specific adaptive process to the TME component. While we classified M5–EMT above as a physiological response to fibroblasts, its dynamics over time also suggested that adaptation to the presence of fibroblasts culminates in a decrease of the EMT module, together with upregulation of interferon response and epithelial senescence programs; rendering this fibroblast-influenced cancer cell EMT tendency biphasic as opposed to linear. Adaptive responses to macrophages included cellular stress responses and metabolic changes-related modules that were elevated over time in the MK line. Specifically, M13–Stress accounted for the integrated stress response and oxidative stress (e.g., *Atf4*, *Mt1*), and M15–Hypoxia was enriched with genes related to poor oxygen stress level responses (e.g., *Bnip3*, *Higd1a*).

The third category of modules reflected cell processes independent of the interactions with macrophages or fibroblasts in the co-cultures, such as cell-cycle-related modules (Fig. 2C, bottom left; and Fig. S4). Recovering this source of gene expression variation was a form of positive control for our identification of modules, as this is an expected form of cellular variation within any population of sustained proliferating cells.

Collectively, the patterns of gene module expression over time provided evidence for both physiological and adaptive changes in response to co-culture with the TME (see Methods for additional module descriptions). Interestingly, the adaptive changes were more specific to the TME cell type than the physiological responses, suggesting a distinct mechanism of adaptation to each TME cell type. While a physiological response is a programmed reaction to the presence of the TME cell type, the adaptive changes may account for a phenotypic change also in the macrophages and fibroblasts.

To functionally test whether the adaptive modules that we identified each reflect a coordinated regulatory program or a set of genes that happen to be co-expressed, we sought to perturb a candidate regulator within a module and assess the resulting transcriptional response in the adapted cells. We focused on the M13–Stress module, which is induced upon co-culture with macrophages, and noted that it contains ATF4, a known regulator of stress response and part of the AP-1 machinery^52^. Inferring that ATF4 may regulate the stress response encompassed by the observed adaptive modules, we used distinct CRISPR-cas9 guides to knock-out (KO) and validate the loss of the ATF4 protein (Fig. S11A). To then test for the effect of ATF4 KO’s in the cancer cells’ transcriptomes, we again set up a 1-week co-culture of MK_36_ and MK_36_ ATF4-KO cell lines with macrophages and processed the lines for scRNA-seq (Fig. 2D and Fig. S14F).

Calculating the expression of the ATF4 gene targets, we found – as expected of a regulator – that their expression was significantly reduced in the KO cells (Fig. 2E,F). With this ATF4 KO line we could now test whether or not the M13–Stress module was a set of genes whose expression is dependent on a common regulation (Fig. 2D). We thus calculated the expression of the M13–Stress module, and found a significant reduction in the KO cell line (Fig. 2G, H, Table S1). In addition, other macrophage-adaptation related modules, M14–Metabolic regulation and M15–Hypoxia (Fig. 2C), were also reduced in expression following ATF4 KO (Fig. 2G). The remaining modules were however unaffected, while some cell-cycle related modules, including M2 and M3, showed increased expression in the ATF4 KO line. The M8-Stress P53 module was also found to be significantly decreased in the ATF4 KO line (Fig. S11B). In summary, the perturbation experiment supported our inference that the M13–Stress module represented a set of genes with a common regulator.

To further test the connection between the module expression and the adaptive phenotypes, we next tested whether the tumor growth advantage that we had observed in the MK adapted lines (Fig. 1C) is disrupted upon perturbation of ATF4. For this we performed an *in vivo* experiment where we injected mice with the MK_36_ ATF4-KO cell line, and compared the resulting tumors with those spawned by MK_36_ V2 control, KL_1_, KL_36_, FK_1_, FK_36_, MK_1_ and MK_36_ lines. We found a significant reduction of tumor volume in the MK_36_ ATF4-KO injected mice, suggesting that the observed gene expression changes in the adapted cell lines relate to important tumor-level growth properties (Fig. S12).

### Recurrence, reversibility and generality of the adaptive and physiological modules

The extended duration of the co-culture experiments (36 weeks) made it possible that the adaptive modules arose through a mutational mechanism. Hence, we queried this possibility using three lines of analyses: recurrence of the adaptive modules, reversibility in the withdrawal of the co-culture, and the degree of clonal selection. We also addressed the generality of our observations regarding physiological and adaptive responses by repeating the co-culture experiment with two additional cancer cell lines, corresponding to ovarian and breast cancer.

If the adaptive modules depended on specific mutations, we would expect substantial variation across independent co-cultures, since each would likely acquire distinct mutations. To test this, we asked whether the gene expression changes were recurrent across independent co-culture experiments. We processed the additional three replicates of the experiment from weeks 4, 12, and 24 for scRNA-Seq analysis, and scored the cells for the modules that captured the significant physiological or adaptive responses (Fig. 3A-C, Fig. S14B). We found a highly conserved pattern of expression across the replicates (Fig. 3B, C). For both the physiological responses (e.g. the M6–CXCL secretion, M12–Secretion, M10–Secretion, and M5–EMT modules) and the adaptive modules (e.g., M4–interferon, M9–Epithelial senescence, M13– Stress, and M15–Hypoxia), the patterns reproduced across biological independent replicates (Fig. 3B, C). For example, M15–Stress (adaptive in macrophages co-culture) showed a consistent increase in all MK replicates, and M9–Epithelial senescence (adaptive in fibroblasts co-culture) increased in all FK replicates (Fig. 3B). The recurrence of the adaptive modules across replicates reflected the robust and non-stochastic nature of this induction, and suggested an epigenetic adaptation, though a convergence of genetic changes was not ruled out.

**Figure 3:**
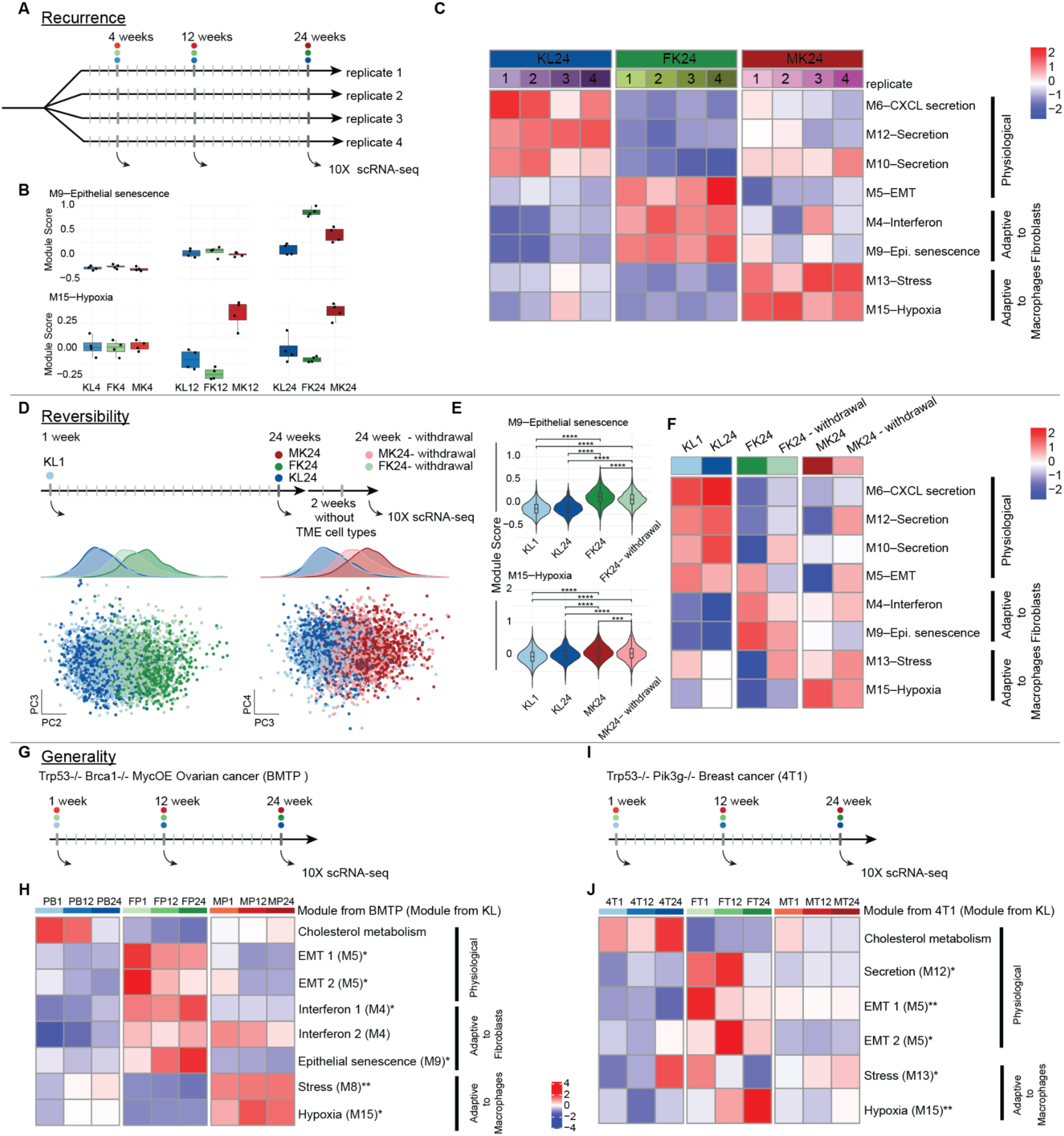
Recurrence, reversibility and generality of the adaptive and physiological modules. **A.** The co-culture experiment was performed in 4 independent biological replicates to test for the recurrence of the observed patterns. **B.** Boxplots indicate scores for module M15–Stress and M9–Epithelial senescence in the four replicates at weeks 4, 12 and 24. **C.** Average scores of the gene modules (Fig. 2) in the replicate co-culture experiments after 24 weeks of co-culture. **D.** Schematic of the withdrawal experiment to test the reversibility of the modules. (top) At 24 weeks of co-culture, we removed the TME cell types and cultured the cancer cells in isolation for 2 weeks. PCA on the module score (bottom), where PC2 and PC3 captured the differences between FK line or MK line and the KL lines, respectively (*P* < 10^-4^). **E** and **F.** Violin plots (**E**) and heatmap (**F**) indicate the extent of stability of the expression of the gene module. Significance was calculated using the t-test. **G** and **I.** Co-culture experiment over 24 weeks using ovarian cancer BMTP (**G**) and breast cancer 4T1 (**I**) cell lines as described for the lung cancer line (Fig. 1). **H** and **J.** Independent Hotspot analysis on ovarian (**H**) and breast (**J**) cancer co-culture data sets captured physiological and adaptive programs. Average scores of these gene modules (Tables 3 and 4) are shown in cells cocultured with fibroblasts (FP or FT, respectively), macrophages (MP or MT, respectively) or in isolation (PB or 4T, respectively). The physiological and adaptive responses are inferred from the module expression level over time and in comparison to the isolated culture, based on the linear model, as was done in Fig. 2. Stars indicate P values for the significance of the change (t-test). Module names were given based on overlapping genes and enriched pathways (Fig. S7 and Methods). In parentheses the corresponding module from Fig. 2 is indicated. * *P* < 0.05, ** *P* < 10^-2^, *** *P* < 10^-3^, **** *P* < 10^-4^.

We next queried for large-scale genomic changes in the co-cultured lines that may be able to explain the expression of the adaptive modules. We did not detect major chromosomal changes when comparing copy number variations across the cell lines over time (Fig. S6), inferred from the scRNA-Seq data^18^ which has been previously used as a robust surrogate for whole exome sequencing^33^. The four independent replicates from the KL, MK, and FK lines showed highly similar CNV profiles at each time point, with a few variabilities in subsets of cells within each replicate, that seemed to acquire changes over time; including a deletion in chromosome 2 (Fig. S6A). Interestingly, the FK cells appeared to accumulate a CNV pattern in all cells over time; in particular, exhibiting changes in chromosome 5, suggesting an accumulative and reproducible change (Fig. S6).

As significant changes in CNVs are known to be associated with cancer cells during EMT^54^, we tested for the possibility that the modules we annotated as adaptive actually corresponded to clonal selection in the culture of existing variation. For this, we clustered the CNV profiles of the MK cells or the FK cells (from all 5 timepoints together), and divided the cells into three CNV-based genetic clones for each group of lines (MK and FK). If the pattern that we observe in the adaptive modules is a result of a clonal selection, we would expect the same clone from the first and 36^th^ weeks to exhibit a similar score relative to the other cells in that timepoint. In contrast, we found that the difference between scores of the adaptive modules followed the same pattern as the overall cells from different timepoints. Yet, we did not find evidence to suggest that a pre-existing clone was present and thus predisposed to the late observed adapted score. Instead we observed a pattern in which all clones showed gradually higher expression of the adaptive modules, indicating that all three clones underwent adaptation, suggesting adaptation in the form of gradual change, rather than clonal selection (Fig. S6C and Fig. S13). We note that while inferCNV did not find evidence for mutations driving adaptation, the possibility remains for smaller-scale variants’ contribution.

We next tested whether induction of the adaptive modules could be reversed upon withdrawal of the co-culture conditions, as a lack of reversibility would be expected from a mutation-dependent change. We cultured the co-cultured lines at the 24-week stage (FK_24_ and MK_24_) for 2 weeks in the absence of their TME cell type (FK_24_-withdrawal and MK_24_-withdrawal, respectively) and asked whether the TME-induced expression remained stable or decayed (Fig. 3D, Fig. S14C). Following these two weeks of culture we processed the cells for scRNA-Seq, scored the cells for the modules and performed PCA on the modules scores (Fig. 3D). We found that PC2 or PC3 captured the differences between conditions, respectively. The FK_24_-withdrawal and MK_24_-withdrawal lines showed significant differences (*P* < 10^-4^) from their FK_24_ and MK_24_ counterparts, and presented an intermediate score between those lines and the ancestral KL lines, suggesting a reverting trend. Examining specific modules, we found that module score patterns across conditions were overall following a similar trend, though with lower scores (Fig. 3E, F). For example, the adaptive M9–Epithelial senescence and M15– Hypoxia still showed the highest scores in FK_24_-withdrawal and MK_24_-withdrawal, respectively, but to a lower extent than the parental co-cultured lines (*P* < 10^-4^ and *P* < 10^-3^, respectively; Fig. 3E, F). The limited stability of the adapted states suggested that the detected modules account for a potentially reversible change. Taken together, the reproducibility, partial stability, and consistent CNV pattern indicated that mutations may not be the sole drivers for the emergence of the adaptive modules.

To test the generality of the appearance of both the adaptive and physiologically induced modules, we asked whether they also emerge in different experimental systems. We considered two additional cancer types by repeating a 24-week version of the co-culture experiment with the BMTP ovarian cancer cell line (*p53^-/-^Brca1^-/-^Myc^OE^-GFP*)^55^ and the 4T1 breast cancer cell line (*Trp53^-/-^; Pik3g^-/-^*, CRL-2539-LUC2). As before, we performed scRNA-Seq on cells from weeks 1, 12 and 24 lines and applied Hotspot analysis to identify gene modules from each of these data sets (Fig. 3G, I, Tables S3, S4 and Fig. S14D). Scoring the cells for the expression of the detected gene modules, we found again that they capture both physiological and adaptive responses. Based on overlapping genes from the gene modules of our lung cancer model (Fig. 2, Fig. S7), we found a common set of cellular responses across cancer types. Specifically, we found that the EMT modules and pattern of expression changes are conserved across the cancer types as a physiological response to the fibroblasts co-culture, followed by downregulation of this program over time as an adaptive response. Also similar to our findings in the lung cancer cell line (KL), we found two interferon modules in the ovarian cancer (BMTP) co-culture, similar to the M4–interferon module in the lung cancer (KL) lines. In addition, in congruence with the KL model, we detected epithelial senescence (M9–Epithelial senescence) as an adaptation to fibroblasts in the ovarian cancer cell line (Fig. 3H). We also identified stress response modules elevated in the co-culture with macrophages in both cell lines (Fig. 3H, J, Fig. S7). The hypoxia signature, found as an adaptation to macrophages in the KL model, was also detected in the ovarian cancer cell line, while in the breast cancer cell line co-culture hypoxia module was found as an adaptive response to fibroblasts, suggesting cancer-type (or driver-type) specific interactions (Fig. 3H, J).

Interestingly, we detected a conserved signature of cholesterol metabolism in the ovarian and breast cancer cell lines. Downregulation of this program appears to reflect a physiological response to the presence of both fibroblasts and macrophages. Such metabolism might be related to hormone-dependent cancers and may be downregulated due to metabolic rewiring in the presence of TME components, or to paracrine lipid supply^56^. Collectively, our observations of both physiological and adaptive responses across the three cancer types provides evidence for a general adaptive process to TME cell types in cancer cells. These results also suggest that the mechanism underlying the adaptive response may be epigenetic in nature.

### Spatial organization of adaptive modules in patient tumor samples

Our results revealed that cancer cells undergo distinct adaptations through the induction of specific gene modules in response to interactions with components of the TME. However, as our results were based exclusively on an *in vitro* co-culture mouse model, we sought to test for their relevance in patient tumors. We therefore employed a spatial transcriptomics (ST) approach, which enabled us to study the *in vivo* spatial relationships between cancer cell states and the composition of the TME in patient tumor samples.

We collected 10 ovarian and 7 breast patient tumors at our medical center and processed them for ST analysis (see Methods). To match the three cancer types used in the co-culture experiments (Figs. 1-3, Fig. S19), we also augmented this dataset with a previously reported lung cancer ST dataset (5 patients)^57^ (Fig. 4A). To first delineate the abundance of major cell types in the TME compartment – fibroblasts, macrophages, endothelial cells, T cells and B cells – we deconvolved the ST data using scRNA-Seq data generated from 3 or 4 of the samples of the ovarian cancer and the breast cancer patients, respectively (see Methods, and^10^). For the lung patient ST data, we determined the cell type annotation based on the original analysis of the data^57^ as well as additional cell type marker expression to generate similar cell type score annotations across samples. As an example of this approach, Figure 4B indicates the abundance of macrophages based on this deconvolution in the OVCA10 ovarian cancer sample. To spatially map the expression of the gene modules found in the mouse cell co-culture experiment, we first matched the co-culture modules to the previously defined patient tumor-based meta programs (MPs) by Gavish et al.^4^ (Fig. 2, Fig. S4). We chose 16 MPs that best matched the modules found in our co-culture catalog and then scored for their expression in all ST^10^ spots (See Methods for MP selection). Figure 4B shows the same ovarian cancer sample scored for the MP5 Stress and the MP1 Cell cycle G2M modules.

**Figure 4.**
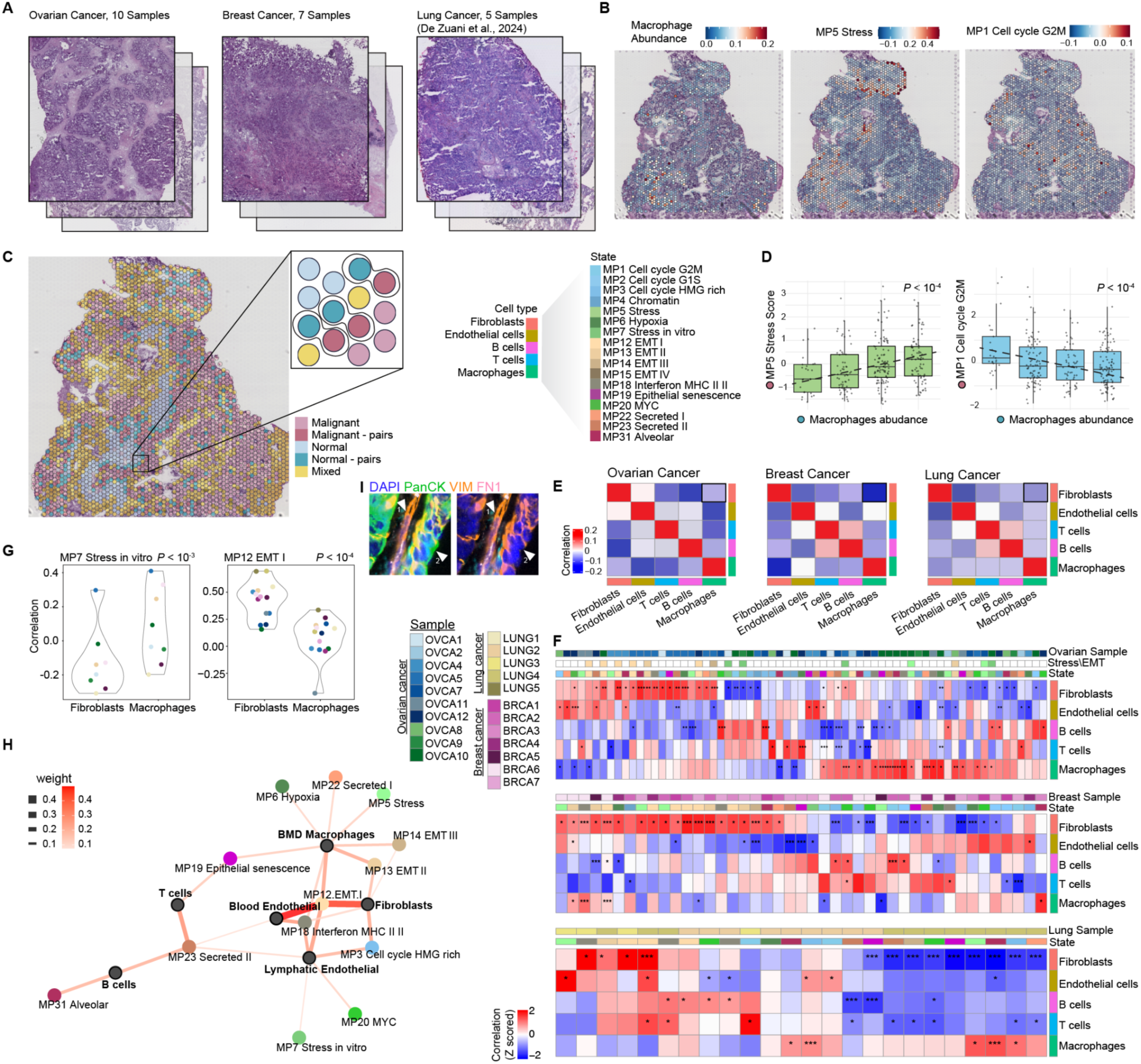
Modules exhibit TME-dependent spatial patterns in patient tumors. **A.** Spatial transcriptomics data set of Ovarian (left), Breast (middle) and Lung (right) cancer patient samples. **B.** Score of Macrophage cell type abundance (left) MP5 Stress (middle) and MP1 Cell cycle G2M (right) meta programs, in ovarian cancer patient sample OVCA10. Cell type annotation is based on deconvolution using previously published single-cell RNA-seq data^10^. **C.** Malignant / Normal / Mixed and border pairs annotation in ovarian cancer patient sample OVCA10 (left). Schematic of the calculation or correlations between cell type score in the Normal-paired and the cancer cell state in the malignant-paired across pairs (right). **D.** Correlation between Macrophage cell type score in the Normal-paired (blue circle) and the MP5 Stress cancer cell state (pink circle, left), and MP1 Cell cycle G2M cancer cell state (pink circle, right) in the malignant-paired across pairs in OVCA10. Significance was calculated using the t-test. **E.** Correlation between the profiles of the major cell types in the cancer cell state colocalization space, in ovarian (left), breast (middle), and Lung (right) cancer spatial analysis. Black boxes mark the macrophages and fibroblasts’ anticorrelation. **F.** Cancer cell states and TME types correlations in all ovarian cancer samples (top), breast cancer samples (middle) and lung cancer samples (bottom). The upper bar indicates the sample, and state (colors as in D). In the ovarian cancer samples (top panel), the middle bar highlights the significant Stress/EMT MPs correlations with macrophages/fibroblasts. Rows and columns are ordered by Zavit^58^. We calculated *P* values in comparison to random pairing of the spots. We filtered the profiles for only the ones that had at least one significant correlation. * *P* < 0.05, ** *P* < 10^-2^, *** *P* < 10^-3^. **G.** Testing for the differences in spatial correlation of MP7 Stress in vitro (left) and MP12 EMT I (right) with macrophages and fibroblasts. Colors represent the samples. We considered only significant values for this comparison. Significance was calculated using the t-test. **H.** Force-directed network graph based on relationships from a single nucleus RNA-seq dataset from 50 lung patients. The graph is annotated for cell types and scored for cell state., where node colors indicate MPs or cell types, and edge thickness and color represent correlation strength. **I.** Multiplexed imaging of the OVCA7 patient sample showing the spatial annotation of fibroblasts (1, positive staining of VIM and FN1) and epithelial cells with EMT markers (2, positive staining of PanCK, VIM and FN1), validating the proximity between fibroblasts and the EMT epithelial cells. Square size 50μm.

To systematically test for co-localizations of expression of the cancer modules and the presence of TME components, we devised a “paired-spot analysis” that examines the TME cell type environments of specific malignant cell states using the ST data. For this we first used inferCNV analysis to annotate the ST spots containing only malignant cells (‘Malignant’), those containing only immune and stromal cells (‘Normal’) and those containing a combination of these (‘Mixed’) (Fig. 4C, Fig. S8, Fig. S19 and Table S8). Next, we delineated pairs of immediately neighboring spots where one is malignant and the other is composed of only normal (non-cancer) cell types. Thus, for every ST sample, we defined pairs of spots with border malignant and immediate neighbor border normal for analysis as pairs. We paired each malignant spot only once to avoid overcounting, and compared the results to random spot pairing (Fig. 4C, Fig. S9, see Methods).

The spot pairings enabled us to study the spatial correlation between cancer cell state expression and TME cell type abundance, avoiding the effect of auto correlation within an ST spot. We computed the correlation between each cancer cell state score in the malignant border spots and the TME cell type abundance in the paired normal spot (Fig. 4C, D). For example, in sample OVCA10, we found significant positive and negative correlations between MP5 Stress malignant cell state and MP1 Cell cycle G2M, respectively, and the presence of macrophages (*P* < 10^-4^ for both, Fig. 4D). When relating the profiles of the TME cell types in terms of their correlations to cancer cell states in all ovarian cancer samples, we found that a major axis of separation is the distinct colocalization of cancer cell states with macrophages and fibroblasts. In other words, macrophage abundance spatially correlates with a specific set of cancer cell states across ovarian cancer samples, while fibroblast abundance correlates with another set (Fig. 4E). Applying this approach to the breast and lung cancer datasets resulted in a similar pattern (Fig. 4E), suggesting that distinct TME cell types are indeed spatially related to different cancer cell states.

To summarize the ST-observed relationships between cancer modules and the TME, we compiled a matrix of the relationships between each of the 16 gene modules and the five TME cell types in the malignant-non-malignant pairs across the 10 ovarian cancer samples (Fig. 4F). Overall, this revealed for each TME cell type a set of correlations and anti-correlations across the specific cancer cell states, indicating a spatial co-existence of these states and TME components (Fig. 4F). We found that stress related MPs, MP7 Stress in vitro and the MP5 Stress, had a spatial correlation with macrophage abundance, while EMT related MPs, MP12 EMT I, MP13 EMT II, and MP14 EMT III, were spatially correlated with fibroblasts, agreeing with the pattern we observed in the co-culture with macrophages (Fig. 4F, significant positive or negative correlations with macrophages or fibroblasts are highlighted in the Stress/EMT bar). To test that the fibroblasts-EMT spatial patterns do not stem from technical difficulty of distinguishing cancer and fibroblast cells in the ST, we used multiplexed imaging (see Methods), and validated with single-cell resolution the proximity between fibroblasts and EMT-like epithelial cells (Fig. 4I).

Extending this analysis to breast and lung cancer samples (Fig. 4F), we found consistent positive spatial correlations between EMT MPs and fibroblasts (Fig. 4F, G). Interestingly, the MP18 Interferon module showed a mixed pattern, with positive correlation with fibroblasts, suggesting that the M4–Interferon module identified in the co-culture analysis (in both lung and ovarian cancer) reflected an interaction with fibroblasts that also occurs *in vivo,* as well as, positive correlation with macrophages. It was previously observed that the interferon signature was typically colocalized with macrophages^10^. While this patient-based spatial analysis provides only a snapshot of the complex interactions and does not allow the separation of the possible physiological and adaptive responses of the cells, they are consistent with the patterns we detected in the co-culture experiment. Moreover, this analysis suggests that our co-culture system may have allowed us to detect interactions that are relevant *in vivo,* but which might be masked by the complexity of the tumor. Notably, stress-related MPs spatially correlated with macrophages, providing additional evidence for the relevance of the modules identified in the co-culture experiments (Fig. 4F, G).

We further tested for an association between expression of the cancer gene modules and cell types of the TME in patient tumors in a single-nuclei transcriptomics dataset from 50 lung adenocarcinoma patient tumors^59^. We hypothesized that samples with a higher cancer cell expression with a particular adaptative module would exhibit a higher fraction of the TME cell types associated with that particular adaptation. To test this, we scored the malignant cells for the expression of the same MPs that represent our modules (See ‘MPs matching to KL co-culture modules’ in Methods), and examined the correlations between the average expression of the MPs in the malignant cells, and the number of cells from each TME cell-type, across patients. We visualized the correlations between cell states (MPs) and cell type abundance in the patients samples, as a force-directed network graph (Fig. 4H). This graph indicated a relationship between fibroblast frequencies and EMT and interferon programs, while macrophages were correlated with stress and hypoxia cancer cell state programs, in agreement with our co-culture findings (Fig. 2C). Interferon and EMT states are also related to macrophages, as well as to endothelial cells, indicating the complexity of these interactions and the challenge to their *in vivo* decoupling, as was also evidenced in our spatial analysis.

Collectively, these analyses provided support for the relevance of the co-culture adaptation modules, as we observed that their expression in patient samples co-localizes with different TME components and generally represents specific possible interactions with the TME cell types. In particular, we found that EMT and stress states preferentially co-localize with fibroblasts and macrophages, respectively (Fig. 4G). We also detected these patterns significantly using another method for spatial transcriptomics (Xenium, Fig. S18). As the interactions within the tumor are multidimensionally complexed in space and time^20,21,34^, our co-culture experiments allowed us to isolate the variables of the system, focusing on specific cell-type interactions and unidirectional adaptation.

### Disruption of a macrophage-adaptive gene module in the cancer cells leads to reduced TAM-like signatures in the co-cultured macrophages

The phenotypic changes that we observed in the co-cultured cancer lines, and which we also captured in the patient data, led us to investigate whether the co-cultured cancer cell lines modulated the state of the TME cell types (macrophages or fibroblasts) with which they were co-cultured (Fig. 5A, Fig. S14E and Fig. S15). As the TME cell types were freshly added weekly, changes in their state as they come into contact with the cancer cells, over the weeks of the experiment, are expected to be a response to changes in the evolving cancer cell lines (Fig. 5A). We thus queried for gene expression changes in the TME cell types after having been co-cultured for four days with the cancer lines of the 1,12, 24, and 36 weeks of co-culture. At four days of co-culture, TME cells were still significantly present in the culture and we were able to include them in the scRNA-seq analysis, and separate them computationally by clustering and cell type markers (Fig. S3) Using principal component analysis (PCA) on the macrophages cultured with cancer cells from different numbers of co-culture weeks, we found associations between their gene expression changes and the number of weeks of the co-culture (Fig. 5B, Fig. S10A, B). PCA analysis allows unbiased quantification of the variability in gene expression within the data, without comparing conditions directly. Studying PC scores across conditions (co-culturing the macrophages with cancer cells from different timepoints) allowed us to test whether these conditions drive substantial changes of gene expression. Detecting those changes between macrophages co-cultured with cancer cells of different timepoints, confirmed that sequencing after four days is sufficient for cellular response.

**Figure 5.**
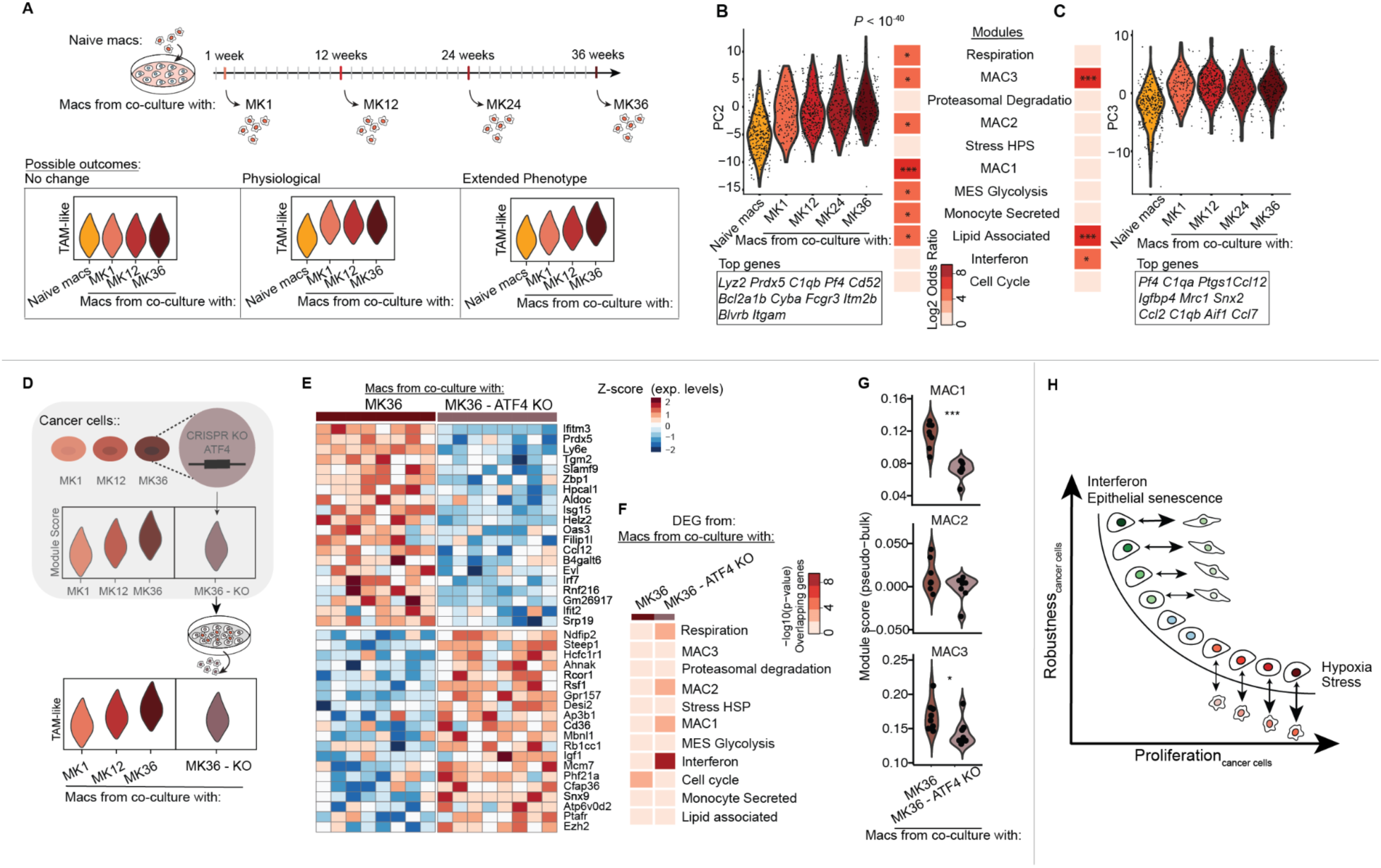
TME pro-tumoral state is driven by cancer cell adaptation to the TME. **A.** Schematics of the experiment. Macrophages were collected from the co-culture, after being co-cultured with MK cells of week 1, 12, 24 and 36. Gene expression of these macrophages can be no different, immediately different or gradually changing compared to naive macrophages. **B, C.** PC2 (**B**) and PC3 (**C**) values for PCA on macrophages co-cultured with the indicated MK cell lines, over 5 time points of the experiment. Cells collected for scRNA-Seq analysis were in co-culture or kept in naive culture for four days. Top contributing genes to each PC are indicated. *P* value calculated for linear change across timepoints. Heatmap indicating the significance of the overlap between the PC genes and TAM signatures. **D.** Schematics of the experiment. Cells were co-cultured with macrophages for 36 weeks, resulting in an increased expression of the M13–Stress module. CRISPR knockout of ATF4, a stress regulator from the M13 module, resulted in a lower expression of M13 in the cancer cells (Fig. 2). Co-culture of these KO cancer cells with macrophages is expected to reduce TAM signature in the macrophages. **E.** Heatmap of differentially expressed genes (DEG) between macrophages collected from the co-cultures with MK_36_ controls and MK_36_ KO cells. **F.** The -log10 (*P* value) of the overlap between the DEG (from **E**) and the TAM programs from Gavish et al. **G.** Violin plots comparing the expression level of Gavish et al. TAM signatures that were found in Fig. 5B.| **H.** Cancer cell adaptation to the microenvironment and its connection to distinct phenotypes in the tumor. Significance was calculated using the t-test. * *P* < 0.05, ** *P* < 10^-2^, *** *P* < 10^-3^, **** *P* < 10^-4^.

We found that Principal component 2 (PC2) captured the order of changes in expression across time (Fig. 5B, Fig. S10A, B). When comparing the set of genes that contribute most to this PC, we found a significant overlap with genes annotated as part of the MAC1 signature – a tumor-associated macrophages (TAM) program linked to inflammation – and the complement pathway activation, as well as other pro-tumoral gene signatures such as MAC2 and lipid association (Fig. 5B, Fig. S10B). The complement pathway is known to be associated with TAMs in lung adenocarcinomas^60^. In contrast, PC3 appeared to account for a physiological response to the co-culture as the major change is solely between the naive macrophages and those co-cultured with cancer cells, regardless of which week of the co-culture was queried (Fig. 5C, Fig. S10B).

Repeating this analysis for fibroblasts co-cultured with the FK lines at the progressing week times, we found that PC2 and PC5 also showed an association with time (Fig. S10 C-D). The set of genes contributing most to PC2 is enriched with previously defined CAF signatures, such as MHC-II machinery-related genes and the complement pathway, including genes such as *Sox4* and *Tagln2*^4^. Together, these changes in the cellular state of the macrophages and fibroblasts suggest a role for adapted cancer cells in eliciting novel states in the TME.

Our experimental design allowed us to distinguish between what are likely physiological TME responses to the presence of cancer cells and those states under the control of the adapting cancer cells. In other words, we are able to distinguish (1) those changes occurring each time the TME cells are co-cultured with cancer cells as an immediate and possibly genetically programmed response to cancer cells and likely anti-tumor, with (2) those occurring progressively each time the TME cells are co-cultured with more adapted cancer cells and are likely pro-tumor. To test this, we investigated the effect of the disrupted expression of an adaptation module in the cancer cells and query for the predicted changes to the TME. To assess the effect of M13–Stress disruption (Fig. 2D-G) on the co-cultured macrophages, we compared the expression of macrophages cultured with MK_36_ with those co-cultured with MK_36_-ATF4-KO cancer cells (Fig. 5D and Fig. S14F). Identifying the differentially expressed genes, we found that genes that are down-regulated in the macrophages co-cultured with the ATF4 KO overlap significantly with TAM-like states, as we had predicted (Fig. 5E,F). This led us to directly examine the expression of the TAM 1-3 module signatures, as they were significantly associated with PC2 and PC3 genes (Fig. 5B-C), capturing the transcriptional change over macrophages co-cultured with more adapted cancer cells. We found that co-culture with MK_36_-ATF4 KO cancer cells led to reduction of their expression in the macrophages (Fig. 5G).

Studying possible ligand–receptor interactions between the MK lines and the macrophages, we further found that the MK_36_ co-culture was enriched for specific pairs relative to the MK1 co-culture. These included interactions associated with macrophage activation and TAM-like states, such as *Csf1*-*Csf1r* and *Ccl3*-*Ccr1*. In comparison with the MK_36_ ATF4-KO co-culture, additional TAM-associated interactions, including *Cx3cl1*-*Cx3cr1*, were enriched in the MK_36_ co-culture (Fig. S17). These results provided further support for the notion that TAM-states were dependent upon the expression of the cancer-cell adaptive M13–Stress module.

## DISCUSSION

In this work we focused on the principles underlying the interactions between the TME and malignant cells, and the resulting cancer cell states, whose importance is highlighted by their influences on tumor progression and response to therapy^2,28,61–63^. A given interaction between a cancer and a non-cancer cell may be driven such that: (1) the cancer-cell state may be a response to the non-cancer cell, (2) the cancer cell may be the director of changes in the non-cancer cell, or (3) the two may co-adapt their responses as a reciprocal interaction. These modes are difficult to observe directly in tumor samples using current approaches, due to the complexity during tumorigenesis and progression in space and time, and so we reasoned that it was necessary to establish indirect means to study these processes. We therefore sought an approach that could isolate specific interactions, while also accounting for the time to enable cancer evolution. These considerations led us to implement an experimental setup designed to isolate specific cancer-TME interactions – outside of the complex tumor ecosystem – whereby the cancer cells were able to evolve while the cell types of the microenvironment were replenished fresh weekly (Fig. 1). While such an experimental setup may miss potential synergistic effects, it does have the power to isolate and carefully study individual effects. As models of the microenvironment, we turned to macrophages and fibroblasts, to represent major cell types within the TME^23,25,26,64–66^. Thus, while co-cultures have been an important tool in many cancer studies^67–69^, our own experimental design was unique as a long-term and single-sided evolution with a transcriptomic read-out.

One first key observation was the delineation of both physiologically- and adaptively-induced gene modules in cancer cells throughout the co-culture experiment. By sampling the co-culture at regular intervals using transcriptomic read-outs, we identified a catalog of modules (Fig. 2), with evidence for these interactions also in patient samples using ST analysis (Fig. 4). We expect that further work will extend this catalog to additional TME cell types and conditions, exploring also, for example, the role of endothelial cells in tumorigenesis^70^. While by design we prevented the TME cells from co-evolving with the cancer cells, it remains plausible that cell types of the TME also adapt or undergo clonal selection, as known to be the case for lymphocytes, and this is also an interesting future direction. Furthermore, our experiment was limited from detecting synergistic module responses of co-culturing with both macrophages and fibroblasts – as these are known to form functional circuits with cancer cells^71^ – as well as other TME combinations.

A second key observation is that the freshly added naive macrophages and fibroblasts adopt increasingly TAM and CAF states in the presence of cancer cells that have been in co-culture with those cell types for a longer time (Fig. 5). When further tested by perturbing a cancer adaptive module, we found that the TAM signature in the macrophage counterpart was reduced. This suggests that as cancer cells adapt to their local TME environment they increase their control over it, to elicit cancer-favorable TME states. Such TME states can thus be considered as an ‘extended phenotype’^72^ of the cancer, as the variants that evolve to establish the TME states are not under selection in the TME cells but rather are present in the cancer cells. As originally proposed by Richard Dawkins, the extended phenotype is premised on the notion that a variant controlling a phenotype in one biological entity (cell or organism) may be under selection remotely in another entity. As an example, the female mosquitoes infected with the *Plasmodium* parasite are significantly more attracted to human breath and odors than uninfected mosquitoes, when the parasites are at a point in their life cycle when they are most infectious to humans^72^. Thus, if we imagine two competing variants of a *Plasmodium* gene, one would be selected for in the parasite population and have its frequency increase if it caused the phenotype in the mosquitoes.

This extends a large body of work already present for cancer cells shaping their surroundings, thus making the TME cancer’s extended phenotype^73^. Evidence for the extended phenotype in cancer comes from patient breast tumors of different subtypes subcutaneously injected in mice, where the obtained tumors exhibit similar microenvironments as in the original tumor^74^. There is also evidence that the genetic driver in lung cancer shapes the TME and affects drug response and treatment outcomes^75^. Another example is the secretion of matrix metalloproteinases (MMPs) that contributes to the breakdown of extracellular matrix components of the parenchymal and vascular basement membranes, normally mechanical barriers to tumor cell migration and invasion, enabling the expansion of cancer. Cancer cells are known to induce this fibroblast MMPs expression by paracrine signaling^76^. We may infer in this case that there is a variant (genetic or epigenetic in nature) in the cancer cells that induces this phenotype, which provides an advantage to the cancer cells that carry this variant. Such benefits may be distributed to the entire cancer population, but spatial proximity may serve as an explanation for the biased benefit, on average, to the cells that carry the variant. Since the criteria for natural selection are met, the fibroblast state can be seen as an extended phenotype of the cancer cell’s variant. As another example, *p53* and *BRAF^V600E^* mutations in cancer cells have been shown, respectively, to induce the M2-like state in macrophages and to lead to CAF states that suppress cytotoxic T cells^77^.

To these anecdotal examples, in this work we define the first catalog of such adaptive responses (Fig. 2). Under this model, heritable variants in cancer cells encode the microenvironment remotely by eliciting selective advantages from the non-cancer cells, and therefore the cancer cells are the ultimate beneficiaries of the interactions with the TME, rather than a model of symbiosis. The lens of the extended phenotype constitutes an attractive approach for generating new insights into cancer TME interactions. More broadly, this approach offers a path towards disentangling the myriad overlapping and adaptive causal relationships occurring in the highly complex tumor microenvironment.

An important consideration regarding the adaptation modules we describe is whether their induction is generally encoded by genetic mutations or cellular plasticity. Interestingly, studying the reversibility, recurrence and clonal relationships, we did not find evidence that the adaptive module inductions could be obviously explained by genetic changes, however we did not exclude the possibility of causal single-nucleotide changes (Fig. 3). Meanwhile, evidence for mutational-independent mechanisms has accumulated for the induction of cancer cell states in terms of their recurrence, plasticity and co-option of existing programs^3,4,10,78–80^. As both genetic mutations and epigenetic changes are stably heritable, they can both be subject to natural selection, and consequently the extended phenotype model is agnostic regarding the mechanistic encodings of variants. If cellular plasticity does play a role, it remains an open question by which mechanisms the cell is able to adaptively find novel interactions without mutations. Similarly to developmental constraints^13,81–83^, answering this question may require the identification of novel avenues and constraints available to cells, particularly once they have accumulated the oncogenic drivers to achieve plastic states.

Finally, we note the possibility that ‘extended phenotype’ adaptations may occur in a cell autonomous fashion, rather than the level of the population. We have recently introduced a possible mechanism for adaptive genome regulation in which a cell’s changing pool of AP-1 transcription factors encodes its cellular plasticity^84^. It is conceivable therefore that as cells are tuning their expression levels in attempts of reducing cellular stress, these effects lead to changes to their micro-environment. In this formulation, the reshaping of the environment is still a result of an evolutionary process – though in this case it is an intracellular process. This process explains our observations that all clones induce the adaptive modules over the weeks of the experiment. In fact, we perturbed here one AP-1 subunit, ATF4, to test for its role in directing module gene expression and observed its effect on the microenvironment (Fig. 5 and Fig. S12).

We propose that considering the tumor through the lens of the extended phenotype amounts to a powerful framework for revealing new principles of cancer biology. The extended phenotype model can help to clarify the underlying logic of interactions between cancer and non-cancer cells and makes testable predictions; in particular that (1) the observed adaptations must provide a fitness advantage to the cancer cells, and (2) the benefit of a state of the non-cancer cell must preferentially feed back to the causal variant in the cancer cell (perhaps by virtue of its spatial proximity). Moreover, it would be interesting to test for cancer’s extended phenotype with modifications of the non-cellular compartments of the tumor microenvironment, such as ECM arrangement and oxygen availability. Finally, a systematic approach similar to the one taken in this study – isolating distinct interactions and examining them in isolation of the vastly complex tumor – can be directed at additional questions including the maintenance of heterogeneity in cancer cells states, the evolution of drug resistance, and the steps to metastasis. New avenues for therapeutic interventions may be revealed by studying how cancer cells adapt to shape their environment and how this imparts them with differential tumorigenic potentials

Finally, our third key observation from the *in vitro* co-culture model system was that cancer cells adapt distinctly to different TME cell types: while the macrophage-adapted cancer cells (MK) induced stress and hypoxia modules, the fibroblast-adapted cancer cells (FK) induced EMT and epithelial senescence modules. This imparted the MK cells with increased tumorigenicity *in vivo,* and the FK cells exhibited enhanced EMT and drug resistance (Fig. 1). These aspects of heterogeneity shed light on the plasticity of states that has been documented on macrophages and fibroblasts^26,85–87^. Furthermore, such a relationship between the adaptations by the cancer cells as they shape their environment, mirrors the recurrent dichotomy in life history known as r/K theory^88–90^, where opposing yet successful strategies are high proliferation and robustness.

Our results suggest that there may also be a proliferation-robustness axis of adaptation as a fundamental tradeoff of cancer cellular states as they are challenged by elements of the microenvironment (Fig. 5H). It would be interesting to search for additional “archetypes” as well as intermediate states among these^91^. Future work is required to test for the mechanism, the fitness, and the role of the stress module of TME-adapted cancer cell states.

As the interactions under study here are complex we reduced the dimensionality – including both the cell cultures and the spatial transcriptomics – requiring further work to understand the full dimensions of the tumor. In particular, our experimental setup carries important limitations to consider. We performed the co-culturing of cells according to the optimal growth conditions of the cancer cells thereby placing the TME cell-types in sub-optimal growth conditions. Moreover, it is likely that our experimental set-up can be optimized such that additional TME phenotypes may be observed with more prolonged co-culture conditions. It is also possible that the experimental design permits the evolution of passage-fitness phenotypes; though in this work we focused on modules specific to particular co-cultures. We note that our implementation of the long term co-culture set-up with its cell passaging, may have led to cell populations that become confluent and arrested on a weekly basis; though we performed all readouts on day four of each cycle, and did not detect a confluency effect at that point. Finally, in this work we did not attempt to elucidate the precise mechanism by which cancer cells communicate with TME-cell types though this is an important line of research for future studies.

## METHODS

### Long-term co-culture experiment

#### Primary fibroblast and macrophage cultures

Primary fibroblasts and macrophages were prepared on a weekly basis. Fibroblasts were isolated from the lungs of C57BL/6 mice perfused with phosphate-buffered saline (PBS). Lung tissue was finely minced and enzymatically digested in a solution containing collagenase D (400 Mandl U/ml; Roche) and DNase I (1 mg/ml; Sigma-Aldrich) at 37 °C with constant shaking.

Following digestion, tissue fragments were cultured in Advanced DMEM supplemented with 1× GlutaMAX (Gibco), 15% fetal bovine serum (FBS), and 1× antibiotic-antimycotic (Gibco), and maintained in a hypoxic incubator (37 °C, 5% CO_2_, 3% O₂) for two weeks. Resulting fibroblasts were then transferred to EMEM supplemented with 15% FBS, non-essential amino acids, sodium pyruvate, and 1× antibiotic-antimycotic, and cultured under normoxic conditions (37 °C, 5% CO₂) for an additional two weeks prior to co-culture experiments. Macrophages were derived from bone marrow cells harvested from the femur and tibia of C57BL/6 mice. Cells were cultured in RPMI 1640 medium supplemented with 10% FBS, 1× antibiotic-antimycotic, and murine macrophage colony-stimulating factor (M-CSF; 20 ng/ml; BioLegend) for 7 days prior to co-culture experiments.

### Cancer cell lines

Murine cancer cell lines used for co-culture included: KL (*KrasG12D*; *Lkb1*−/− lung adenocarcinoma, derived from a genetically engineered mouse model and provided by the laboratory of K.-K. Wong) and transfected with LARRY-GFP library, BMTP (*p53*−/−; *Brca1*−/−; *MycOE-GFP* high-grade serous ovarian carcinoma organoid line, provided by the laboratory of B. Neel) ^93^, and 4T1 (triple-negative breast cancer, obtained from ATCC). Single-cell clones from each line were established and used in co-culture assays. The KL clone was isolated after the LARRY infection, to ensure a single clone was isolated. KL and 4T1 cells were cultured in RPMI 1640 supplemented with 10% FBS, 1× GlutaMAX, and 1× antibiotic-antimycotic. BMTP cells were maintained in high-glucose DMEM supplemented with 3% FBS, 1× GlutaMAX, 1× sodium pyruvate, 1× ITS supplement (Gibco), 1× antibiotic-antimycotic, and murine EGF (50 ng/ml).

### Co-culture experiment

For co-culture assays, 1×10⁴ KL or 4T1 cells, or 5×10³ BMTP cells, were seeded in 6-well plates either alone, with 1×10⁵ fibroblasts, or with 5×10⁵ macrophages in their respective growth media supplemented with M-CSF (20 ng/ml). Fibroblast cell number (1 x 10^5/well in 6-well plate) following primary culture isolation protocol as optimal growth (protocol by the Wong lab). To approximately match over all cell density in the plate, and account for the macrophage high abundance in lung cancer, a macrophage: fibroblast ratio was set at 5:1. The KL cell number (1x10^4^ /well in 6-well plate), was set to allow cell growth and reach confluency after 7 days of culture, to allow less frequent cell passaging. The long-term co-culture was performed in 4 independent replicates for the KL line, and in 2 independent replicates for 4T1 and BMTP lines. Each cancer line was expanded from a single isolated clone. After 7 days, co-cultures were subjected to a two-step dissociation: cells were first treated with mild TrypLE Express (Gibco) to detach fibroblasts, followed by 0.25% trypsin to detach remaining cancer cells. Recovered cancer cells (1×10⁴) were mixed and reseeded into fresh wells with newly prepared 1×10⁵ fibroblasts, 5×10⁵ macrophages, or alone. At the time of passaging, after 7 days of culture and with the described passaging protocol, the reseeded cells were dominantly cancer cells only.

This weekly co-culture cycle was repeated for 24 weeks for BMTP and 4T1, and for 36 weeks for KL cells. Cells from the cultures at weeks 4, 12, 24 and 36 were frozen for future experiments.

### Proliferation assay

KL, MK, and FK cell lines from weeks 1, 12 and 36, were plated at a concentration of 200 cells per well in a 48 well plate. Macrophages or fibroblasts added at 1:50 or 1:10 ratios. Each condition was plated in 5 technical replicates. Plates were placed in an Incucyte incubator for culture and imaging for 4 days. The analysis was done based on 9*9 imaging, every 8 hours. Cell count was done by the Incucyte software algorithm and was normalized to cell count on the first round.

### Cell preparation for single-cell RNA-seq

Cells were collected at the designated week of co-culture. For additional experiments, cells were frozen down, then thawed and re-cultured for two weeks with fibroblasts, macrophages or in isolation, and collected at day 4 of co-culture, on the designated week (or on day 4 of the second week of re-culture, if cells were thawed). Cancer cells in isolation (KL lines) were washed with PBS, and incubated with 0.25% Trypsin for 20 minutes at 37 °C. Cells were then collected with 1:1 fresh media, followed by washing and suspension. For the FK cells, the cells underwent the two-step dissociation as mentioned above. Briefly, cells were first treated with mild TrypLE Express (Gibco) to detach fibroblasts, followed by Trypsin for 20 minutes at 37 °C.to detach remaining cancer cells. MK cells underwent a similar two-step process, where first cells were treated with trypsin to dissociate the cancer cells, and then the remaining macrophages were scraped and collected separately. Cancer cells, fibroblasts and macrophages were counted separately. For the main long-term co-culture experiment described in figure 2, fibroblasts or macrophages were mixed with their parental cancer cell line, at a 70:30 ratio, demultiplexed together, and recovered both cell types from each well. For downstream analysis the cell types were separated bioinformatically based on clustering, and annotated based on cell hashing and marker expression.

### 10x Genomics library preparation and sequencing

We used the 10x Genomics workflow for scRNA-seq on cells collected for the comparison between conditions and timepoints. Cell suspensions from different conditions were labelled using the 10x Genomics 3′ CellPlex kit Set A according to the manufacturer’s instructions. Cells were pooled and loaded into the 10x Genomics Chromium Controller. Cell loading for droplet-based scRNA-seq and library preparation was performed using the 10x Genomics Chromium platform with the Chromium Single Cell 3.0 Library & Gel Bead Kit v3.1 following the manufacturer’s instructions. Libraries were diluted to 2 nM, and 75 bp paired-end sequencing was performed using the Illumina NextSeq 500. 150–200 million paired reads were generated for each library. Raw reads were processed with the CellRanger v3.1 or v.7.0.0 (10x Genomics) pipeline (default parameters) with the mm10 genome references. Demultiplexing of the corresponding CMOs was performed using CellRanger v.7.0.0 with 0.8 as assignment confidence. For the replicates, single-cell RNA-seq analysis (Fig. 3), cells from all 4 replicates, of the 3 conditions (KL, MK, FK) at a given time point, were demultiplexed, pooled and divided into two 10x reactions, which were loaded simultaneously to the 10x and were sequenced together. Schematic of all co-culture experiments leading to separate 10X sequencing described in Fig. S14A and Tables S7 and S8.

### Single-cell data analysis

#### Filtering

Data was loaded using the ‘Read10X’ function, followed by Seurat R package^94^ v4.3.0 pipeline. Briefly, for each cell, the number of transcripts, genes expressed, and the proportion of transcripts derived from mitochondrial and ribosomal genes were determined. For each dataset, cells with transcript numbers lower than 700 or higher than 2× s.d., with less than 500 genes detected and with higher than 20% of transcripts corresponding to mitochondrial genes were excluded from further analysis. Genes expressed in less than 1% of the cells in each dataset were removed.

### Cell clustering and differential gene expression analysis

Dimensionality reduction, clustering and differential expression were performed using the Seurat ^94^ R package v4.3.0. Highly variable genes were identified using the Seurat FindVariableGenes function (default parameters) and these were used to compute the PCA following UMAP using the top five to ten principal components (cutoffs determined by the elbow method). For cell type separation, clustering was performed using the FindClusters function using the selected principal components with resolution set to 0.2. Differential gene expression across treatments was performed using the FindMarkers function using the MAST test (logfc.threshold=0.25, min.pct=0.2, min.cells.genes=10).

Differential expression analysis – at the bulk level – between only the cancer cells across the treatments revealed down-regulation of seven *CXCL* genes in both co-cultures relative to the TME-free KL line (Fig. S3D). We found that the macrophage co-cultured MK lines showed up-regulation of the DNA-binding inhibitors, *Id2*, *Id3*, and *Far1*, that have been associated with aggressive cancer phenotypes and unfavorable prognoses^95^. In contrast, In the fibroblasts co-cultured FK lines, we observed an overexpression of *Ctla2a*, *Sparc*, and *Fn1*, suggesting an epithelial-to-mesenchymal transition-like (EMT) state. *Sparc* overexpression was previously shown to activate fibroblasts in endometrial cancer^96^ (Fig. S3D).

### Hotspot analysis

A graph-based method to find modules agnostic of condition or time point was implemented according to DeTomaso et al.^41^, Hotspot method, in Python, as described here https://github.com/yoseflab/hotspot. Comparison with human datasets and published gene modules was done using the Ritan R package^97^ with the molecular signatures database (MSigDB) hallmarks^43,44^, following conversion of the gene modules to human analogues using the biomaRt R package^98^.

### Module scoring

To assign the score for the expression level of the Hotspot modules, we used AddScoreModule by Seurat. For cell cycle scoring, we used the Seurat function ‘CellCycleScoring’ to assign cells to G1, S, and G2/M phases. For averaged analyses, we used AverageExpression by Seurat.

### Linear Model calculation

Linear model coefficients and p-values were calculated using the ‘lm’ function in R. The calculation was done at the level of FK vs KL or MK vs KL, and with the addition of time.

### Module descriptions

We classified the modules into three broad categories, physiological, adaptive, and independent of co-culture conditions. In the physiological response category, M6–CXCL secretion, M12– Secretion, and M10–Secretion were related to chemokine secretion. Specifically, M6–CXCL secretion was enriched with Cxcl genes (e.g., *Cxcl1*, *Cxcl2*), which are known to be secreted to elicit immune cell recruitment^45–47^. These *Cxcl* ligands are *Cxcr2* ligands, and their overexpression is associated with cell adhesion and migration^48^. M10–Secretion included secreted molecules (e.g, *Lif*, *Epha2*) together with cell surface adhesion molecules (e.g., *Cd44*, *Itga3*), and M12–Secretion with secreted molecules related to matrix proteins (e.g., *Anxa2*, *Fbln2*). This suggests that the physiological response to both fibroblasts and macrophages is characterized by a broad decrease in expression of *Cxcl* ligands and matrix reorganization. M5– EMT was strongly associated with EMT (see Results).

The adaptive category of modules included gene expression increase in the cancer cell co-cultured with fibroblasts (FK lines) over time; M4–Interferon which included genes related to interferon response and MHC class II (e.g., *Ifitm1*, *Ifitm3*); M9–Epithelial senescence, an epithelial senescence program (*Slpi*, *C3*, *Eomes*); and M11–Myc targets, a program involving Myc targets (e.g., *Dkc1*, *Nop56*, *Kmt5a*). Cellular stress responses and metabolic changes-related modules were elevated over time in the MK line. M13–Stress appeared to account for the integrated stress response and oxidative stress (e.g., *Atf4*, *Mt1*). M14–Metabolic regulation was enriched with genes related to cell migration and growth (e.g., *Cdh6*, *Marcks*) and metabolic regulation (e.g., *Ak3*, *Ogt*), and M15–Hypoxia was enriched with genes related to hypoxia (e.g., *Bnip3*, *Higd1a*).

The third category of modules reflected cell processes independent of the interactions with macrophages or fibroblasts in the co-cultures, such as cell-cycle-related modules (Fig. 2C bottom left and Fig. S5). Modules M1 and M3 corresponded to the G2M and G1/S states of the cell cycle, respectively (e.g., *Top2a*, *Mki67*, *Pcna*, *E2f1*). Modules M2 and M7 were enriched with chromatin remodeling genes (e.g., *Suz12*, *Smarca5*; *Kmt2a*, *Nfat5*). Additionally, M8 captured DNA damage-related stress genes (e.g., *Trp53inp1*, *Fosb*, Fig. S5A). Recovering this source of gene expression variation was a form of positive control for our identification of modules of co-expressed genes, as this is an expected form of cellular variation within any population of proliferating cells.

An independent Hotspot analysis was done for the ovarian and breast cancer cell lines (Fig. 3) and the modules detected were named based on their overlap with the KL based modules (see results). In the ovarian cancer cell line, ‘Interferon 1’ was enriched with MHC related genes such as *CD74*, *H2 - Ea*, *H2-Q6*, and ‘Interferon 2’, which included the *Ifim6* gene. Both modules show high overlapping genes with the *MP Interferon MHC II-I* program by Gavish et al. ‘Interferon 1’ also overlaps with *MP Interferon MHC II-II,* and Interferon 2’ shows overlap with MP *Stress* programs by Gavish et al.^4^. Both interferon modules in the ovarian cancer cell line co-culture experiment showed an increase in co-culture with fibroblasts similar to that in the KL co-culture, but also a gradual decrease in the co-culture with macrophages suggesting different paths of adaptation to each TME cell type. Moreover, a stress response module captures a physiological response to macrophages, and highly overlaps with the M8–Stress P53 module from the KL co-culture. In the breast cancer line it overlapped with the M13–Stress module from the KL co-culture, and similarly to the pattern in the KL co-culture, it captured the pattern of adaptation to macrophages (Fig. 3H, J, Fig. S8). In addition, two EMT related modules were found in the ovarian cancer co-culture experiment, where EMT1 overlaps with EMT II and EMT 2 overlaps with EMT III programs by Gavish et al. and represent ECN remodeling with genes such as *Fn1, Lamc2 and Sparc,* and epithelial differentiation with Krt19 and Wfdc2, *respectively*. In the breast cancer co-culture *experiment*, also two EMT modules were found. *Both overlap with* Gavish et al. EMT I and II, but EMT 1 overlaps also with the Gavish MES program and indicates a more mesenchymal state.

### inferCNV analysis for the coculture cell lines

The inferCNV R package v.1.6.0^99^ R was used to account for possible genomic alterations. To compare the CNV profiles of the 4 independent experiments, included in Figure 3, the original clone (KL_1_) reference was added to each of the experiments. For the main experiment, with KL, FK and MK lines from weeks 1, 4, 12, 24 and 36 of co-culture that were used in Figs. 1, 2, and 5, KL_1_ cells served as the reference set of cells. CNV profiles were grouped by cell line (week and condition).

To further analyse the differences in module score between inferCNV subclones, the profiles of the main experiment (Fig. S6B), the profiles of each treatment (KL, MK, FK) from all timepoints were allowed to cluster resulting in a hierarchical clustering tree. To allow enough cells from all timepoints in each clone, for module score comparison, the tree was cut such that it resulted in three CNV-based clones. The module score of the cells in each subclone was calculated and compared across timepoints within each subclone.

### Reversibility experiment

To assess the reversibility of the adaptive transcriptional states, cancer cells were collected after 24 weeks of co-culture with either fibroblasts or macrophages (FK24 and MK24, respectively). The TME cells were then removed, and the cancer cells were maintained in culture in the absence of fibroblasts or macrophages for an additional 2 weeks (labeled as FK24 - withdrawal and MK24 - withdrawal). Cells were subsequently collected for scRNA-seq, in addition to the parental co-culture cell lines-KL24, FK24 and MK24. All cells were scored for the previously identified gene expression modules. Module scores were compared between the parental co-cultured cells, the corresponding 2-week withdrawal cultures, and the ancestral KL cells to assess the extent to which the adaptive transcriptional programs were maintained or reverted following removal of the TME stimulus.

#### Ligand-receptor interaction inference

Ligand-receptor pairs were obtained from the CellPhoneDB^100^ database downloaded from GitHub https://github.com/ventolab/cellphonedb-data/. Ligands were defined based on expression in cancer cells and the corresponding receptors based on expression in the co-cultured macrophages. For each ligand-receptor pair, an expression-based interaction score was calculated as the product of the mean normalized ligand expression across cancer cells and the mean normalized receptor expression across macrophages. Interaction scores were compared between MK36 and the corresponding reference co-culture (MK1 or MK36 ATF4-KO) and expressed as log2 fold change. The top 40 positively enriched ligand-receptor pairs were used for visualization.

### Spatial data collection and processing

From the ten ovarian cancer tumors (OVCA1, OVCA2, OVCA3, OVCA4, OVCA5, OVCA7, OVCA9, OVCA10, OVCA11, OVCA12), and the 7 breast cancer tumor samples (BRCA1-7) two to three pieces were embedded in optimal cutting temperature compound (OCT) by placing them cut side down into a plastic mold. The OCT-filled mold was then snap-frozen in chilled isopentane and stored at −80 °C until use. Cryosections were then cut at 10 µm thickness and mounted onto Visium arrays. Tissue optimization and library preparation were performed according to the manufacturer’s instructions, with 12 min of permeabilization. Libraries were sequenced on an Illumina NextSeq and aligned using the Visium SpaceRanger pipeline. As a quality control step, spots with fewer than 700 UMIs or more than 30% mitochondrial or ribosomal reads were filtered out. The Seurat single-cell transformation was used to normalize and center the data, and to identify variable genes.

### ST basic analysis with Seurat and CARD deconvolution

To deconvolve the ST samples at the cell type level, we first utilized previously published single-cell RNA-Seq data from samples: OVCA NYU1 (OVCA5), OVCA NYU2, OVCA NYU3, and BRCA1, BRCA2, BRCA3, BRCA4^10^ and preprocessed using Seurat^10,101^. Samples were merged by sample (OV, BR). The Seurat single-cell transformation was used to normalize and center the data, and to identify variable genes. Data was then integrated using Harmony^102^ .. Cell types within the single-cell data reference, including malignant and non-malignant annotations, was based on the previously described approach by Barkley et al, in “Cell type identification” and “Malignant cell identification” Method sections^3^. Briefly, cell types were assigned based on marker gene expression: B cells (CD79A, CD79B, IGKC, CD14), epithelial cells (KRT7, KRT8, KRT19), stromal cells/fibroblasts (COL1A1, COL1A2, BGN, DCN, VIM), endothelial cells (CLDN5, PECAM1, VWF), T cells (CD3D, CD3E, CD3G), and myeloid cells (CLEC9A, CD1C, CD1A, CD1C, CD1A, CLEC9A) which were further divided into monocytes (CD68, LYZ, AIF1) and macrophages (CD14, FCGR3A, FCGR1A, CD68, TFRC, CCR5). Cell annotations of the single cell data were used for CARD ST deconvolution^103^.

### Malignant spot annotation

ST spots were classified as ‘Malignant’, ‘Normal’, or ‘Mixed’ using a two-step approach. First, spots were clustered within each sample using k-means clustering (k = 3). For initial annotation of the spots into the three categories, the expression of a cancer-associated marker gene (WFDC2^104^ for ovarian cancer and MGP^105^ for breast cancer, as was previously described^10^), was used to classify the clusters to high, intermediate, and low marker expression, together the CARD deconvolution derived cell-type abundances. ‘Mixed’ spots represent spatial regions in which malignant and non-malignant cells are present within the same spot. Second, RNA-based copy-number variation profiles were inferred using inferCNV (v1.6.0)^99^ to identify and refine malignant CNV patterns. CNV clusters were evaluated together with cell-type marker expression and CARD-derived cell-type abundances. inferCNV was iteratively rerun using annotated ‘Normal’ spots as the reference to refine the assignments. Spots with ambiguous CNV profiles were classified as ‘Mixed’. Final annotations were spatially validated against the corresponding H&E images (Fig. S19). The lung cancer samples annotation was based on the authors cell type annotation^57^.

### Matching meta programs (MPs) to the KL co-culture modules

As Gavish et al.^4^ used a comprehensive approach for consensus modules across different cancers, we reasoned that this is an appropriate conversion from the mouse co-culture to modules with relevance to patient tumors. Notably, the conversion need not be one-to-one, as genes are not unique to specific meta-program (MP) and the analysis was performed with a different approach from our own module definitions; thus more than one MPs may match one of our modules and vice versa. Starting with 26 MPs with assigned functions and general occurrence across cell types, we selected those that highly overlapped with our modules (Fig. S5). For example, the M4–Interferon module identified as adaptation to fibroblasts overlapped significantly with (1) MP18 Interferon MHCII, (2) MP23 Secreted II, and (3) MP13 EMTII. Our M9–Epithelial senescence’ module matched (1) MP19 Epithelial senescence and (2) MP31 Alveolar. The M14–Metabolic regulation was matched with (1) MP5 Stress and M15–Hypoxia matched (1) MP6 Hypoxia, (2) MP14 EMTIII, and (3) MP5 Stress. Overall this resulted in the selection of 16 MPs which we scored for expression in all ST spots (See methods and^10^). Figure 4B shows the same ovarian cancer sample scored for the MP5 Stress module. While we found that module M13–Stress overlapped strongly in terms of genes with MP7 Stress in, the MP5 Stress’ may represent adaptation to macrophages nonetheless that was found in our co-culture, as the overlapping genes with these modules include *DDIT3*, *NR4A1* and *NFKBIZ*, that are related to stress response and inflammation. Moreover, this MP highly overlaps with M8–Stress P53 from the co-culture data, noteworthy that M8 is lowly expressed in the MK cells, but still shows a linear increase over time, indicating adaptation rather than a physiological response, only in the MK cells.

### Spatial transcriptomics ‘Malignant’ - ’Normal’ pairs annotation

For the spatial analysis, each ‘Malignant’ spot was paired with its immediate neighbor ‘Normal’ spot, if available. The neighbors’ calculation was done using Euclidean distance on the pixel coordinates of each sample. Immediate neighbors of each spot were found by the shortest distance between spots. If a ‘Malignant’ spot had more than one immediate ‘Normal’ neighbor, a paired ‘Normal’ was chosen randomly. Once a ‘Normal’ spot was paired, it was removed from the potential neighboring list to avoid over-counting and the pair was fixed for the rest of the analysis. ‘Malignant’ spots that did not have a ‘Normal’ immediate neighbor were discarded from the analysis. Each ‘Malignant’ spot was paired at most once in the analysis. As a negative control, we also created sets of randomly paired spots, relative to which we were able to compute the significance of the observed correlations. This approach allowed spatial correlation to be dependent on information from two separate proximal spots, and avoid over interpretation caused by autocorrelation.

### Module score and correlation calculation in ST

Spatial transcriptomic spots from each sample were scored for the expression of the Gavish et al.^4^ meta-program gene signatures using the ‘AddModuleScore’ function in Seurat. Correlation coefficient was calculated for every sample separately, taking the cancer cell state score (Gavish et al., meta-program) value of each ‘Malignant-paired’ spot, and the TME cell type abundance (from the deconvolution) for its paired ‘Normal-paired’ spot. Calculating these correlations resulted in a 16 (cancer cell states) by 5 (TME cell types) matrix for every sample. For ovarian cancer, the final matrix of all the samples together was 260 (26 cancer cell states from 10 samples) over 5 TME cell types. For clustering of these profiles, we used Zavit ^58^.

### Patient tumor multiplexed imaging (CODEX)

Fresh frozen samples (OVCA1, OVCA7, OVCA10, BRCA1, BRCA2) were cryosectioned at 10 µm thickness and mounted onto a glass coverslip coated with polylysine. The tissue was stained and the CODEX multicycle reaction performed according to CODEX user manual

Revision C. Briefly, the sample coverslip was placed on Drierite beads for 5 min, then incubated in acetone for 10 min and set in a humidity chamber for 2 min. The sample was hydrated in CODEX Hydration Buffer twice for 2 min, fixed in 1.6% paraformaldehyde in CODEX Hydration Buffer for 10 min, then washed in CODEX Hydration Buffer twice for 2 min. The sample was then equilibrated in CODEX Staining Buffer for 30 min then stained with a barcoded antibody cocktail in CODEX Blocking Buffer for 3 h in a humidity chamber. Antibodies comprising the antibody cocktail: Pan-Cytokeratin, VIM, FN1. The sample was washed three times for 2 min in CODEX Staining Buffer, fixed in 1.6% paraformaldehyde in CODEX Storage Buffer for 10 min, and then washed three times in 1× PBS. The sample was incubated in 4 °C methanol for 5 min, washed three times in 1× PBS and then fixed using the CODEX Final Fixative Reagent Solution for 20 minutes in a humidity chamber. The sample was washed three times in PBS and then stored in CODEX Storage Buffer for 3 days until imaging. A commercial Akoya CODEX instrument and a Keyence BZ-X800 microscope with a ×20 Nikon PlanApo numerical aperture 0.75 objective were used to treat and image tissue using complementary fluorescent reporters. The protocols from Akoya CODEX user manual revision C were followed. The samples were imaged and processed in one CODEX run. Images were processed using Halo.

### Xenium-based spatial analysis

We assembled a multi-cohort Xenium dataset comprising 3 available 10x Genomics Xenium Prime 5K datasets representing breast, lung and ovarian cancers: three independent datasets representing these cancer types^106^ and six lung cancer datasets^107^. We retained the malignant and tumor-microenvironment cell annotations provided with each source dataset. For each focal malignant cell, we detected all neighboring cells within a 30-μm radius and calculated the local cancer-cell fraction after excluding the focal cell itself. Malignant cells with a local cancer-cell fraction below 60% were defined as border malignant cells and retained for downstream analysis. Each cell’s transcriptome was smoothed using the average of itself and its 4 nearest neighbors. Malignant-cell expression programs were quantified using the manuscript-derived meta-program gene sets using genes represented on the Xenium Prime 5K panel. Within each sample, local fibroblast and macrophage abundances were calculated as their respective fractions among all non-malignant cells in the same 30-μm neighborhood. Sample-level spatial associations were then quantified using Pearson correlations between the smoothed malignant-cell program scores and the corresponding local fibroblast or macrophage fractions. Stress programs were evaluated against local macrophage abundance, whereas EMT programs were evaluated against local fibroblast abundance. Within each sample, we performed a linear regression to test for a positive trend in the module score and the studied cell type fraction.

Samples with a positive slope and P<0.05 were retained for visualization (Fig. S18). For each retained sample, the corresponding fibroblast and macrophage correlations were displayed and compared in a paired analysis. Sensitivity analyses varying the local cancer-fraction threshold, neighborhood radius and malignant-cell score-smoothing scheme produced qualitatively consistent patterns.

### Cell state-cell type correlations

We obtained single-nuclei RNA sequencing data from 50 lung adenocarcinoma patients^59^. We annotated malignant and non-malignant cell types, based on inferCNV and gene markers expression. For each patient we counted the number of cells for each cell type (bone marrow derived (BMD) macrophages, T cells, B cells, lymphatic endothelial, blood endothelial, and fibroblasts), and for the malignant population, we scored for the expression of the MPs (Fig. 4C), and took the average expression of each MP. We visualized the correlations between cell states’ average level (MPs) and the cell type numbers across patients, as a network graph using the ‘*igraph’* R package. Nodes represent either cell states (MPs) or cell types. Edges represent positive correlations above a threshold of 0.1, with edge thickness proportional to the correlation coefficient and edge color reflecting correlation significance. The network layout was computed using a force-directed algorithm (Fruchterman–Reingold), optimizing node positioning to minimize edge crossing and enhance interpretability.

### *In vivo* tumor formation and measurement

All animal studies were performed under protocols approved by the Institutional Animal Care and Use Committee (IACUC) at NYU Langone Health. Five-week-old albino C57BL/6 mice (B6(Cg)-Tyrc-2J/J; The Jackson Laboratory) were acclimated for one week prior to experimentation. KL, FK and MK cells from different timepoints along the co-culture were first co-cultured in isolation, with fibroblasts or with macrophages, respectively, for a week, and were then expanded without fibroblasts or macrophages in 150-mm culture dishes for 5 days. Cells (2×10⁶) were resuspended in sterile PBS and subcutaneously injected into both flanks of recipient mice using a 27-gauge needle. Tumor growth was monitored biweekly by caliper measurements, and animal body weight was recorded concurrently. Mice were euthanized by CO₂ asphyxiation when tumors reached humane endpoints (tumor ulceration), at which point tumors were excised and measured. Two independent experiments showed similar results (Fig. 1C and Fig. S2). An additional independent experiment was performed comparing tumors spawned by the KL_1_, FK_1_, FK_36_, MK_1_, MK_12_, MK_24_, MK_36_ as well as KL_36_, MK_36_ ATF4-KO and MK_36_ V2 control lines (Fig. S12).

### Pathology assessment of tumors based on H&E stating

For hematoxylin and eosin staining, frozen sections were fixed in cold methanol at –20 °C for 30 minutes, followed by a 1-minute incubation in isopropanol. After air-drying, the slides were incubated with hematoxylin for 3 minutes at room temperature and then rinsed with water. A 2-minute bluing step was performed using Bluing Buffer, after which eosin counterstaining was carried out at room temperature for 1 minute. Finally, the slides were dehydrated, cleared, and mounted for imaging. Pathology assessment of representative tumors of each condition, based on H&E stating, was done to grade and assess necrotic levels of the tumors. Intertumoral stroma, fibroblastic reaction, as well as intramural inflammation appear to be more abundant in FK1, FK36, MK1, MK12, MK24, MK36 in comparison to KL1. The assessment of the lung adenocarcinoma histologic grade was done according to College of the American Pathologists D. Histopathologic Grade. The predominant histological pattern of non-mucinous adenocarcinomas is associated with prognosis with lepidic-predominant tumors having the best prognosis, acinar and papillary-predominant tumors having an intermediate prognosis, and solid-predominant and micropapillary-predominant tumors having the worst prognosis. The IASLC Histopathologic Grading Scheme for Non-mucinous Lung Adenocarcinoma was recommended for resected early-stage non-mucinous lung adenocarcinoma: Grade 1 (G1): Well-differentiated (lepidic-predominant with no or < 20% high-grade pattern) Grade 2 (G2): Moderately differentiated (acinar or papillary-predominant with no or < 20% high-grade pattern) Grade 3 (G3): Poorly differentiated (any tumor with ≥ 20% high-grade pattern (that is, solid, micropapillary, cribriform, or complex glandular pattern).

A complex glandular pattern is defined by fused glands or single cells infiltrating in a desmoplastic stroma.

### Cell viability assay

Five hundred cells were seeded in tissue culture-treated 96-well plates and incubated overnight at 37 °C. Media was supplemented with eight concentrations of drug, in 6 technical replicates, and incubated in 37 °C for 3 days. Cell viability was determined using a Cell Counting Kit-8 (Cat #ALX-850-039-0100; Enzo Life Sciences), according to the manufacturer’s protocol.

Luminescence was measured at 450 nm, 4 h after adding CCK-8 reagent to the cells using a FlexStation 3 multi-mode microplate reader, according to the manufacturer’s instructions. The concentrations measured for the G12D Kras inhibitor (MRTX1133) and for the Chemotherapy drug (Paclitaxel) are 10, 1, 0.1, 0.05, 0.025, 0.01, 0.001, 0 uM and 10, 1. 0.5, 0.25, 0.1, 0.01, 0.001, 0 uM, respectively

### Cell line CRISPR knock-out generation

gRNAs for gene knock-out were designed using the Benchling software and cloned into Cas9 lentiCRISPR v2 Blast (Addgene, Cat #83480) vector.

Guide sequence: sgAtf4 g1 - CCGCCAACACTTCGCTGTTC, g2 AGATGAGCTCTTGACCACGT. For lentivirus production, HEK-293T cells were co-transfected with Cas9 lentiCRISPR v2 Blast along with the packaging plasmids psPAX2 (Addgene, Cat #12260) and pMD2.G (RRID: Addgene_12259) using Lipofectamine 3000 (Invitrogen, Cat #L3000008), following the manufacturer’s protocol. Cell culture supernatants containing viral particles were collected 72 h later and purified through 0.45-μm filters (Corning, Cat #431225) to eliminate cellular debris. Subsequently, viruses were used to infect various cell lines via a 1-hour spin-infection at 600 × g at 32 °C in the presence of 8 μg/mL polybrene (Sigma-Aldrich, Cat #TR-1003) and subsequent blasticidin (Invivogen, Cat # ant-bl-05) selection 24 hours post-infection.

### Immunoblotting

Cell pellets were lysed using RIPA buffer (Thermo Fisher Scientific, Cat #89900,) with protease and phosphatase inhibitor cocktail (Thermo Fisher Scientific, Cat #78440;). Proteins were quantified using a Pierce™ BCA Protein Assay Kit (Thermo Fisher Scientific, Cat #23225). Subsequently, equal quantities of total protein from each sample were mixed with 6x reducing Laemmli SDS sample buffer (Boston BioProducts Inc., Cat #BP-111RR) and distilled water, and then boiled at 95 °C for 5 minutes to achieve protein denaturation. Subsequently, the samples were loaded and electrophoresed on 4–20% Criterion TGX Precast Midi Protein gels (Bio-Rad, Cat #5671094) and then transferred onto nitrocellulose membranes for immunoblotting. The membranes were incubated overnight at 4 °C with primary antibodies targeting ATF4(Abcam Cat# ab270980) and β-actin (Sigma-Aldrich Cat# A5441, RRID:AB_476744). For detection, IRDye 800CW donkey anti-rabbit IgG (LI-COR Biosciences Cat# 926-32213, RRID:AB_621848) and IRDye 680RD donkey anti-mouse IgG (LI-COR Biosciences Cat# 926-68072, RRID:AB_10953628) secondary antibodies were incubated for 1 h at room temperature, washed three times with PBS containing Tween 0.01%, and visualized using an Odyssey detection system (LI-COR Biosciences).

## Data and Code Availability

The single cell RNA-seq and spatial transcriptomics results were deposited to the NCBI GEO database with the accession code GSE328449 for single cell RNA sequencing, and GSE206165 for spatial transcriptomics data. Code used in this study is available at https://github.com/MaayanPour/Ext-Phen-scripts.

## Acknowledgements

The authors thank L. Keren, N. Erez, T. Papagiannakopoulos and the members of the Yanai laboratory for helpful discussions; the laboratory of B. Neel for providing cell lines and the laboratory of M. Maurano for assistance with sequencing. We are grateful to NYU Langone Experimental Pathology Laboratory for their assistance with sectioning, staining and imaging services. This work was supported by the following NIH grants: This work was supported by the following US National Institutes of Health grants: R01 CA296978, U01 CA260432, R01 LM013522, U54 CA263001 and R21 CA264361.

